# Selective abundance of miR-290-295 in the adult *Substantia Nigra* dopamine neurons is neuroprotective via preservation of protein synthesis

**DOI:** 10.1101/2025.08.05.668587

**Authors:** Zixuan Li, Yang Xu, Nicola Murgia, Rongpu Liu, Ece Tasci, Nikolay Kovzel, Dick San Ng, Haixia Jiang, Yu Liu, Xuejia Kang, Donglei Sun, Andrii Domanskyi, Wenjie Zhang, Ilya A. Vinnikov

## Abstract

Locomotor, reward and other critical functions of the body are regulated by the ventral midbrain, with the central role played by dopamine (DA) neurons. The function of these cells from the early development to maturity is critically dependent on the orchestrated expression of coding and non-coding genes. For example, in the stem cells the miR-290-295 cluster constitutes the majority of expressed microRNAs and is critical for stemness in rodents. During development towards various terminally differentiated lineages, such as neurons, the cells typically switch off transcription of these stem cell-specific microRNAs. Here we report that within the adult *Substantia Nigra pars compacta* (SNpc), the miR-290-295 cluster is exclusively expressed in DA neurons (SN^DA^), preventing the locomotor deficits and maintaining an adequate expression of proteins involved in DA biogenesis, such as tyrosine hydroxylase (TH), dopa decarboxylase (DDC) and DA transporter (DAT). Importantly, a global knock-out of the miR-290-295 cluster leads to decreased numbers of SN^DA^ neurons in adult mice. Using *in vitro* and *in vivo* DA cell-specific loss-of-function models, we demonstrated that miR-292a-3p, the most abundant microRNA in this cluster, directly targets PTEN, a phosphatase antagonizing the neuroprotective phosphatidylinositol-4,5-bisphosphate 3-kinase (PI3K)-AKT-mechanistic target of rapamycin kinase (mTOR) pathways regulating translation initiation. Accordingly, when labelled with the click chemistry-compatible methionine analogue L-azidohomoalanine, miR-290-295 cluster-deficient SN^DA^ neurons revealed a drastic impairment of protein synthesis, which is critical for DA biogenesis. Our surprising findings demonstrate for the first time a selective expression of stem cell-specific and stemness-promoting microRNAs in a distinct population of mature neurons to maintain their physiological functions in the adulthood, suggesting that similar epigenetic disinhibition mechanisms may be also critical for other terminally differentiated cells across species.

**Graphical abstract:** 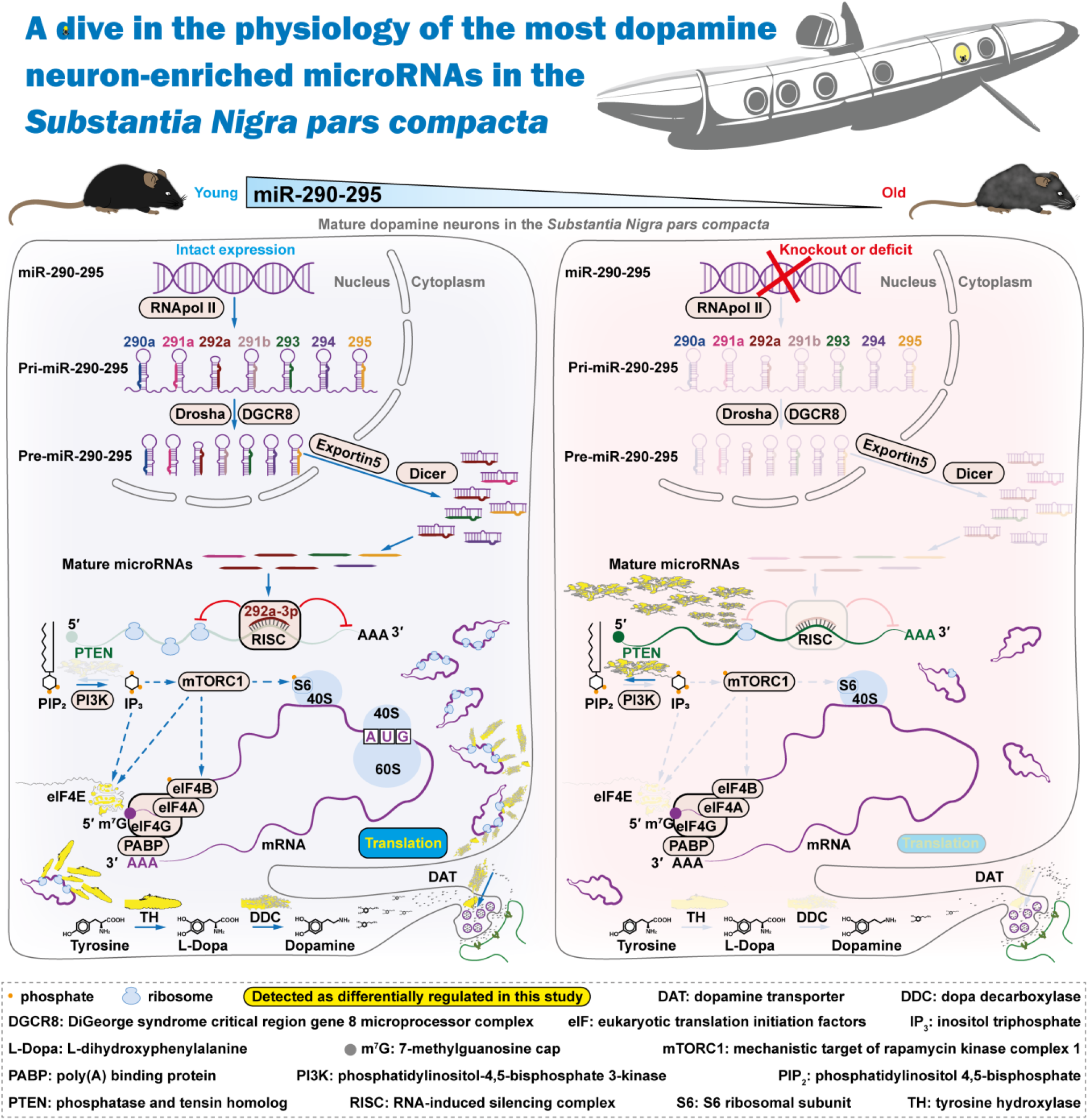

## INTRODUCTION

The dopamine (DA) system is crucial for reward, cognitive control, emotion, motivation, appetitive behavior and sexual arousal locomotor and other functions [1–4], with its alteration leading to neurodegenerative diseases, such as Parkinson’s disease (PD) [5]. The development and physiology of DA neurons are critically dependent on the orchestrated expression of microRNAs, small non-coding RNAs capable of degrading transcripts and blocking their translation [6], which are especially relevant for local regulation at the presynaptic terminals.

Since the early embryonic development, pluripotency is strongly dependent on high expression of the miR-290-295 cluster and other stemness-promoting genes [7, 8]. Constituting more than 60% of all microRNAs expressed in the embryonic stem cells, this cluster promotes pluripotency, proliferation and senescence, while inhibiting apoptosis [8–13]. Not surprisingly, both during embryonic development and regeneration in the adulthood, the cells lose these stemness-promoting microRNAs upon differentiation [8]. Compared to the mature oocyte, after a modest decline by 17% and 33% in the zygote and the two-cell stage blastomers, respectively, expression of miR-290-295 gradually rises in four and eight-cell stage blastomers [14], morula and the inner cell mass of the blastocyst, but not the differentiating trophectoderm which at this stage reveals a drastic down-regulation of the cluster [7]. In gastrula and later developmental stages, a further decline of the cluster expression occurs in differentiating tissues [15] with trace levels still detected in neural precursors [8, 16], developing neural tube, heart and other actively proliferating tissues on the mouse embryonic days 8.5-18.5 (E8.5-18.5) [17, 18]. Interestingly, miR-290-295 and their paralogs from the cluster miR-302 sharing the same seed region are critical for neural tube closure by preventing premature differentiation of neural epithelium into precursors and neurons [10, 19]. Accordingly, in the adulthood, the expression of the miR-290-295 cluster declines upon differentiation [18, 20], with miR-292a-3p still detected in the limbs of E14.5 [20], but not adult mice [21–25], miR-294-3p expressed in the developing heart during prenatal stages, but lost in the neonate and adult heart [26]. In the adult central nervous system, the miR-290-295 cluster is abundant in the stem cell niches, such as the hippocampus [27] or the subventricular zone [16]. Using highly sensitive analytical techniques, only trace quantities of the cluster microRNAs can be detected in mature γ-aminobutyric acid (GABA), glutamatergic neurons or Purkinje cells [28], total brain [29] or midbrain [30], but not striatal [31] isolates.

Many microRNAs regulate both neuronal maturation during development and physiology of DA and other neurons in the adulthood. Indeed, brain-specific microRNAs, such as miR-9-5p and miR-124-3p, promote neurogenesis and differentiation during the development, as well as synaptic plasticity in mature neurons [32–34], with the latter microRNA maintaining DA neurons by counteracting apoptosis and autophagy [35–37]. Declining in multiple brain regions of PD patients, including the *Substantia Nigra* [38, 39], miR-34b/c-5p promotes cell cycle exit and induces dopaminergic differentiation [40]. During embryonic development, miR-128-3p increases the average dendritic spine length of NSCs [41], enhances neurogenesis, inhibits proliferation and promotes differentiation of NPCs into neurons by suppressing the pericentriolar material 1 expression [42]. Another microRNA declining in the brain tissues of PD patients, miR-7-5p is highly abundant in and critical for maturation of DA and other neurons [43–45]. miR-186-3p promotes apoptosis of DA neurons and disrupting DA biogenesis via inhibition of insulin-like growth factor 1 receptor (IGF1R) and DA transporter (DAT), respectively [30]. While remaining neuron-specific in the adult brain, this microRNA promotes DA neuronal survival in MPTP neurotoxicity mouse model by reciprocal down-regulation of axis inhibition protein 1 (AXIN1) and up-regulation of excitatory amino acid transporter 4 (EAAT4) expression, despite impairing autophagy and increasing neuronal vulnerability to α-synuclein toxicity [30, 46]. Abundant in the *Substantia Nigra pars compacta* (SNpc) DA neurons (SN^DA^), miR-133b-3p regulates the maturation and physiology of these cells by targeting paired like homeodomain 3 (Pitx3) [47–49]. Thus, similar to coding genes, such as nuclear receptor subfamily 4 group A member 2 (NR4A2, or NURR1), forkhead box A1/2 (FoxA1/2), paired-like homeodomain transcription factor 3 (PITX3), engrailed homeobox 1 (EN1/2), LIM homeobox transcription factor 1 α/β (LMX1a/b) and orthodenticle homeobox 2 (OTX2) [50–54], microRNAs often play a critical role both during embryonic or postnatal development of the nervous system and in the maintenance of DA and other mature neurons.

Interestingly, DA neurons in the ventral tegmental area (VTA) of two-week-old rat pups express miR-290 [55], but no studies reported expression of any of the cluster member in DA neurons of adult rodents. Indeed, stem cell-specific genes promoting stemness, such as miR-290-295, are typically not detected in the brain [56–58] or midbrain [59, 60], unless highly enriched from large amounts of isolated material [29, 30]. This study was focused on mature aimed to identify the most mature DA neuron-enriched microRNAs in SNpc and determine their functions in the adulthood.

## RESULTS

### Within the mature SNpc, the miR-290-295 cluster is selectively expressed in DA neurons

First, we performed high throughput profiling of SNpc aiming to identify microRNAs that are highly enriched in DA neurons, i.e. that are scarcely or not at all expressed in the neighboring cells within this region. For that, tamoxifen-induced Dat^Cre-ERT2^:Dicer1^fl/fl^ or Control littermates [4, 61] were stereotaxically injected into the striatum with retrobeads to fluorescently retrogradely label their DA neuron bodies within the ventral midbrain. 4 or 6 wk after tamoxifen injection, SNpc regions were dissected from coronally sliced freshly isolated brains using an ellipse-shaped tip tissue puncher with its opening encircling the shape of the SN^DA^ neuronal groups and analyzed by TaqMan quantitative reverse transcription polymerase chain reaction (qRT-PCR) microRNA array (**Figure 1A** and **Tables S1-S2**). Selective knockout of Dicer in SN^DA^ neurons is expected to cause the loss of all Dicer-dependent mature microRNAs in these neurons, but not in the surrounding neuronal and non-neuronal cells. Thus, we hypothesized that down-regulated mature microRNAs would be the ones highly enriched in SN^DA^ neurons. Surprisingly, we detected the stem cell-specific miR-290-295 cluster as the most down-regulated microRNAs in adult mice after Dicer deletion from DA neurons, compared to Control mice (**Figure 1B-C**). These results indicate that i). intact SN^DA^ neurons express these microRNAs and ii). they are highly enriched or exclusive to these cells within the heterogenous SNpc tissue. Sharing a highly conserved seed regions (**Figure 1D**), all microRNAs in the miR-290-295 cluster are expressed from the same promoter [62] and functionally promote stemness [11, 12] and cell survival [11], while also preventing apoptosis [11, 63, 64]. Our data indicate that several members of this cluster are as abundant in the adult SN^DA^ of control mice as a well-studied miR-133b-3p [47, 65, 66] or miR-124-3p, which co-expresses with miR-124-5p, one of the most abundant microRNAs in mature dopaminergic and other types of neurons [35, 36, 67–71] (**Figure 1C**).

**Figure 1.**
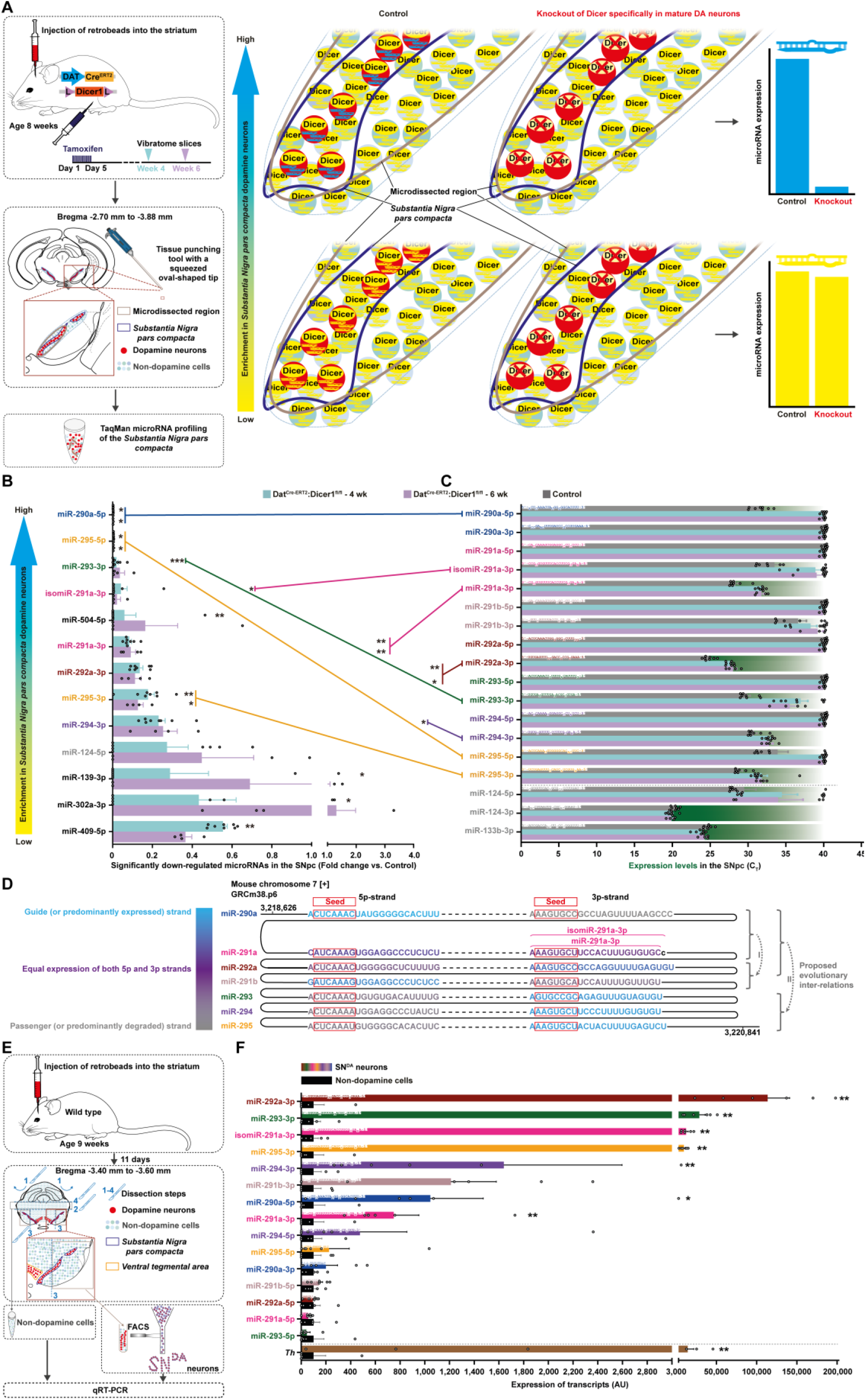
miR-290-295 are the most dopamine neuron-specific microRNAs in the *Substantia Nigra pars compacta*. (**A**) Experimental design to induce Dicer knockout in mature dopamine (DA) neurons by intraperitoneal (i.p.) tamoxifen or sunflower oil injections of 8-wk-old Dat^Cre-ERT2^:Dicer1^fl/fl^ mice of both sexes, followed by stereotaxic injections of retrobeads into the striatum and microRNA profiling of the tissue puncher-isolated *Substantia Nigra pars compacta* (SNpc) tissues. Biological replicates were as follows: Control, 5 females and 3 males; Dat^Cre-ERT2^Dicer1^fl/fl^-4wk, 3 females and 5 males; Dat^Cre-ERT2^Dicer1^fl/fl^-6wk, 3 females and 1 male. (**B-C**) Significantly down-regulated microRNAs in Dat^Cre-ERT2^:Dicer1^fl/fl^ compared to Control mice (**B**) and the expression of the miR-290-295 cluster and other DA neuron-abundant microRNAs (**C**) in SNpc of Dat^Cre-ERT2^:Dicer1^fl/fl^ 4 and 6 wk after recombination or Control mice (n = 8, 4 and 8, respectively). (**D**) Schematic illustration of the genomic context and phylogenetic conservation of miR-290-295 based on the GRCm38.p6 genome assembly. Seed regions are outlined with red color. (**E**) Experimental design of fluorescence-activated cell sorting (FACS) of SNpc DA (SN^DA^) neurons. Red retrobeads were injected into the striatum of 9-week-old wild type male mice, and samples were harvested for fluorescence-activated cell sorting (FACS) 11 days later. Biological replicates were as follows: Non-dopaminergic cells, 5 10-11-week-old wild type males; SN^DA^ neurons, 6 biological replicates generated by pooling samples from a total of 19 10-11-week-old wild type males (5 replicates were each pooled from 3 mice, and the final replicate was pooled from the remaining 4 mice). (**F**) The relative expression levels of indicated microRNAs normalized with *U6* in SN^DA^ neurons versus non-dopaminergic cells (n = 6 and 5, respectively). Error bars represent standard error of means (SEM). *, *p* < 0.05; **, *p* < 0.01; ***, *p* < 0.001; ****, *p* < 0.0001 as analyzed by Kruskal-Wallis test with Dunn’s multiple comparisons test versus the Control group (**B**), unpaired two-tailed Mann Whitney test or unpaired two-tailed Student’s t-test versus the non-dopaminergic cells (**F**). See the details about ages, sexes, FACS gating strategy and statistical methods in the **Source data**.

To validate these surprising findings, we next assessed the levels of the miR-290-295 cluster specifically in SN^DA^ neurons of adult mice. For that, we injected retrobeads into adult mice, dissected *Substantia Nigra* and collected positive for the tracer by fluorescence-activated cell sorting (FACS). *Th* was used to validate the enrichment of SN^DA^ neurons in the samples. Consistent with our previous results, we observed a strong enrichment of the miR-290-295 cluster within SN^DA^ neurons (**Figure 1E-F** and **Tables S1-S2**), which was further confirmed for selected microRNAs in a complementary experiment employing FACS-sorted DAT-positive SN^DA^ neurons (**Figure S1A-D**).

### Expression of the miR-290-295 cluster in SN^DA^ neurons decreases with aging

To further confirm these data, we stereotaxically injected retrobeads into striata of wild type mice, with the aim to selectively microdissect SN^DA^ neuron bodies by laser-assisted technique (**Figure S1E** and **Table S1**). In addition to the young adult mice, we also used 23.5-month-old animals to analyze the expression of the cluster microRNAs during aging. Interestingly, despite the lower obtained amount of total RNA in comparison to FACS-assisted experiments, SN^DA^ neurons revealed expression of miR-291a-3p, miR-293-3p, miR-294-3p, miR-295-3p and the most abundant miR-292a-3p at the levels comparable to miR-124-3p and miR-133b-3p, with the latter three of these cluster microRNAs declining upon aging (**Figure S1F** and **Table S2**).

### Knock-out of miR-290-295 leads to reduced SN^DA^ neuronal numbers in adult mice

Next, to explore the functions of these microRNAs in midbrain DA neurons, we assessed their numbers in mice with a global knockout of one or both alleles of miR-290-295 [10]. Strikingly, at the age of 35 wk, one out of the five homozygous mutants revealed tail suspension swings [72], posture asymmetry and locomotor deficits (**Video S1**), characteristic for parkinsonism-like neurodegenerative phenotype. Moreover, we detected an allelic dosage-dependent decrease of SN^DA^ neurons and the density of their projections in mice with both heterozygous and homozygous knockout of the cluster, as assessed by immunostaining for the critical proteins in DA biogenesis: tyrosine hydroxylase (TH), dopa decarboxylase (DDC) and DAT (**Figure 2A-I**). Interestingly, DA neurons in the VTA only revealed a declining tendency (**Figure S2A**), suggesting that the cluster is especially critical for the development of SN^DA^ neurons and, possibly, also their survival in the adulthood. This is in line with their well-studied higher vulnerability [73–76] stemming from a vicious cycle combining toxic DA metabolism, mitochondrial complex dysfunction and distinct physiological constraints [77]. Since replenishing of stem cell populations and their asymmetric division is critical for proper embryonic development, it might not be surprising that maturation of the central nervous system is affected by the lack of stemness-promoting microRNAs, which suggests a critical function of this cluster during development, with a specific bias to the SN^DA^ lineage. Either an alternative or an additional interpretation of the above data suggests a physiological role of these microRNAs in maintaining SN^DA^ neurons in the adulthood.

**Figure 2.**
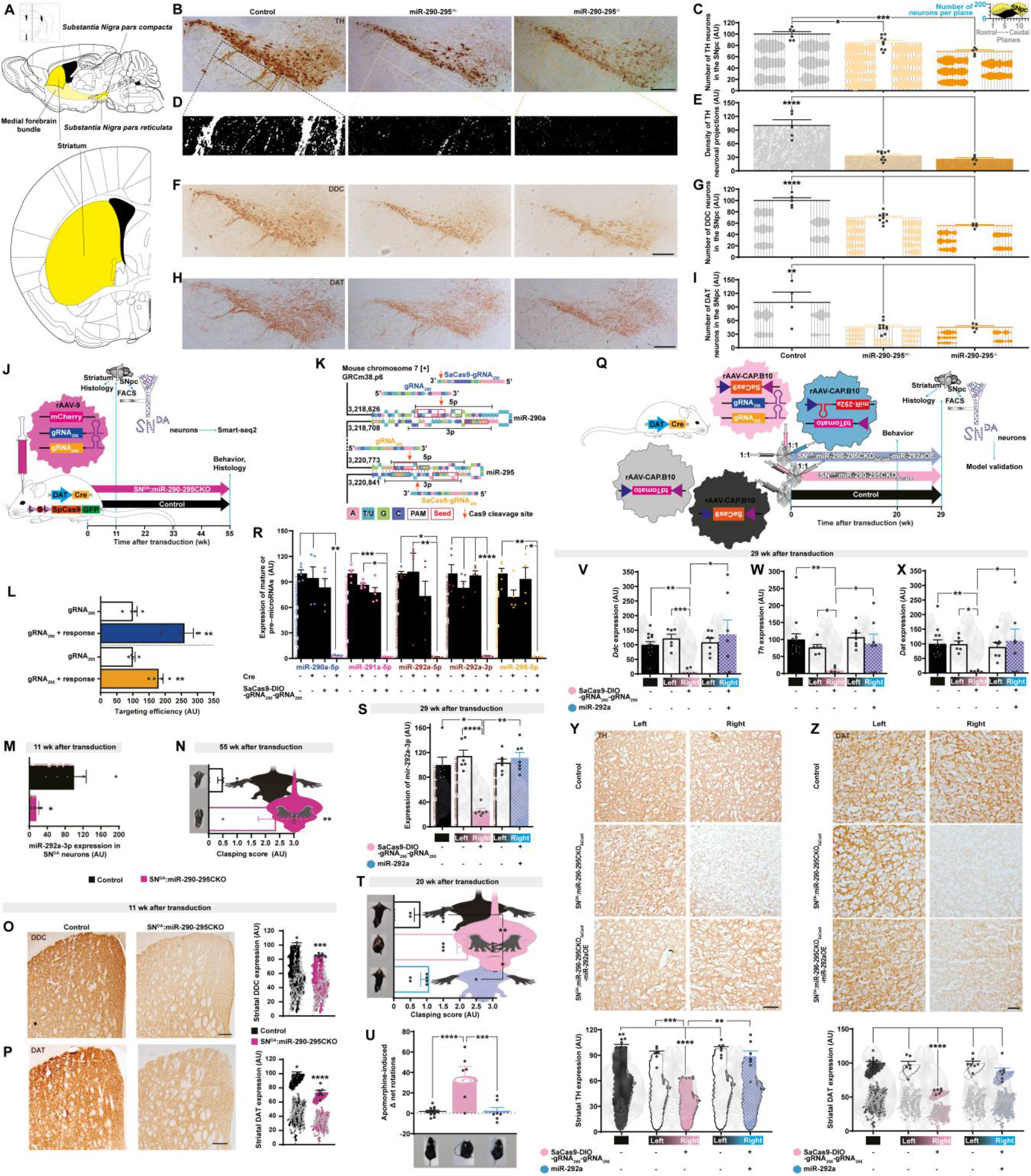
miR-290-295 is critical for SN^DA^ neuron physiology. (**A**) Anatomic location of the SNpc, the medial forebrain bundle and the striatum. (**B-C**) Representative microphotographs (**B**) and quantification of the numbers of SN^DA^ neurons (**C**) immunostained for tyrosine hydroxylase (TH) in miR-290-295^+/-^, miR-290-295^-/-^ and Control mice (n = 9, 5 and 6, respectively). (**D-E**) Visualization (**D**) and quantification of the area of TH-immunostained projections in the *substantia nigra pars reticulata* (**E**) in miR-290-295^+/-^, miR-290-295^-/-^ and Control mice (n = 9, 5 and 5, respectively). (**F-I**) Representative microphotographs (**F,H**) and quantification of the numbers of SN^DA^ neurons (**G,I**) immunostained for dopa decarboxylase (DDC, **F,G**, n = 10, 5 and 5) and dopamine transporter (DAT, **H,I**, n = 10, 5 and 4) in miR-290-295^+/-^, miR-290-295^-/-^ and Control mice, respectively. (**J**) The scheme of the experiment to delete miR-290-295 specifically in mature SN^DA^ neurons. rAAV-9-gRNA_290/295_ was stereotaxically injected into SNpc of mice of 7-14-month-old DAT^Cre^:SpCas9-GFP mice (6 females and 7 males). For the Control group, rAAV-9-empty was injected into SNpc of 7-16-month-old DAT^Cre^:SpCas9-GFP or wild type mice (5 female and 12 males). (**K**) gRNA design for SpCas9 and SaCas9 targeting the miR-290-295 cluster. The scheme illustrates gRNA target sites, PAM motifs, and genomic coordinates selected for SpCas9 (PAM = NGG) and SaCas9 (PAM = NNGRRT) to enable efficient deletion of the miR-290-295 cluster. (**L**) gRNA_290/295_ on-target efficiency analysis (**L,** n = 4) and knock-out validation in FACS-isolated SN^DA^ neurons from SN^DA^:miR-290-295CKO or Control mice (**M**, n = 4 and 5, respectively). (**N**) Hindlimb clasping test score in SN^DA^:miR-290-295CKO and Control mice (n = 4 and 6, respectively) 55 wk after transduction. (**O-P**) Representative microphotographs and quantification of striatal immunostaining for DDC (**O**) and DAT (**P**) in SN^DA^:miR-290-295CKO and Control mice (n = 7 and 5, respectively) 11 wk after transduction. (**Q**) The scheme of the experiment to delete miR-290-295 or supplied with miR-292a specifically in mature SN^DA^ neurons in SaCas9 system. For the SN^DA^:miR-290-295CKO_SaCas9_ group, rAAV-CAP.B10-DIO-SaCas9-gRNA_290/295_ and rAAV-CAP.B10-tdTomato were stereotaxically injected into the right SNpc, while the rAAV-CAP.B10-empty and rAAV-CAP.B10-tdTomato were stereotaxically injected into the left SNpc of 6 *Slc6a3^em(Cre)^* male mice aged 4-5 months. For the SN^DA^:miR-290-295CKO_SaCas9_-miR-292aOE group, rAAV-CAP.B10-DIO-SaCas9-gRNA_290/295_ and rAAV-CAP.B10-miR-292a-tdTomato were stereotaxically injected into the right SNpc, while the rAAV-CAP.B10-empty and rAAV-CAP.B10-tdTomato were stereotaxically injected into the left SNpc of 7 *Slc6a3^em(Cre)^*male mice aged 4-5 months. For the Control group, rAAV-CAP.B10-empty and rAAV-CAP.B10-tdTomato were stereotaxically injected into the both sides of SNpc of 13 *Slc6a3^em(Cre)^* male mice aged 3-5 months. (**R**) *in vitro* validation of the SaCas9-gRNAs design (n = 5 per group except for the SaCas9-DIO-gRNA_290_-gRNA_295_ transfected group when checking miR-295-5p, n = 4). (**S**) Knock-out and overexpressed validation in FACS-isolated SN^DA^ neurons from SN^DA^:miR-290-295CKO_SaCas9_, SN^DA^:miR-290-295CKO_SaCas9_-miR-292aOE and Control mice (n = 6, 7 and 6, respectively) 29 wk after transduction. (**T**) Hindlimb clasping test score in SN^DA^:miR-290-295CKO_SaCas9_, SN^DA^:miR-290-295CKO_SaCas9_-miR-292aOE and Control mice (n = 6, 7 and 13, respectively) 20 wk after transduction. (**U**) Apomorphine-induced Δ net rotations in SN^DA^:miR-290-295CKO_SaCas9_, SN^DA^:miR-290-295CKO_SaCas9_-miR-292aOE and Control mice (n = 6, 7 and 13, respectively) 20 wk after transduction. Δ net rotations = number of full 360° contraversive - number of full 360° ipsiversive rotations. (**V-X**) Expression of mRNAs encoding DDC (**V**), TH (**W**), and DAT (**X**) in SN^DA^:miR-290-295CKO_SaCas9_, SN^DA^:miR-290-295CKO_SaCas9_-miR-292aOE and Control mice (n = 6, 7 and 13, respectively) normalized to glyceraldehyde-3-phosphate dehydrogenase (*Gapdh*) 29 wk after transduction. (**Y-Z**) Representative microphotographs and quantification of striatal immunostaining for TH (**Y**) and DAT (**Z**) in SN^DA^:miR-290-295CKO_SaCas9_, SN^DA^:miR-290-295CKO_SaCas9_-miR-292aOE and Control mice (n = 6, 7 and 13, respectively) 29 wk after transduction. Error bars represent SEM. *, *p* < 0.05; **, *p* < 0.01; ***, *p* < 0.001; ****, *p* < 0.0001 as analyzed by one-way ANOVA followed by post-hoc Holm-Šídák’s test (**C,E,G,I,U,Y,Z**), unpaired two-tailed Student’s t-test (**L-P**), Kruskal-Wallis test with Dunn’s multiple comparisons test (**R,S,T,V-X**), or paired two-way ANOVA followed by post-hoc Holm-Šídák’s test (**S,V-X**). See the details about ages, sexes, FACS gating strategy and statistical methods in the **Source data**. Scale bars: 200 μm.

### Loss of the miR-290-295 cluster in mature SN^DA^ neurons leads to early onset striatal projection impairments

Next, we sought to prove the latter hypothesis in aging, when SN^DA^ neurons become especially vulnerable. We constructed a model with *in vivo* CRISPR/SpCas9-dependent conditional knock-out of the cluster in mature SN^DA^ neurons (SN^DA^:miR-290-295CKO) of at least 4-month-old mice. First, we designed, validated and subcloned two miR-290-295-spanning guide RNAs (gRNAs) optimized for on-/off-target activities into adeno-associated viral vector (rAAV) as described previously (**Figure 2J-L** and **Tables S2-S3**) [78]. The latter was then stereotaxically injected into the SNpc of adult Dat^Cre^:SpCas9-GFP mice to delete the miR-290-295 cluster specifically in mature SN^DA^ neurons (**Figure 2M** and **Table S1**). Interestingly, these mice revealed a late onset hindlimb clasping (**Figure 2N** and **Video S2**), but no decline of striatal DA (**Figure S2B**), SN^DA^ neuronal numbers or projection density in the *Substantia Nigra pars reticulata* up to 55 wk after transduction (**Figure S2C-J**). Interestingly, striatal expression of TH, DDC and DAT in SN^DA^:miR-290-295CKO was significantly down-regulated (**Figure S2K-R**), which was apparent for DDC and DAT as early as 11 wk after transduction (**Figure 2O-P**). These results suggest that the miR-290-295 cluster is critical to preserve the synthesis of TH, DDC and DAT in the terminals of SN^DA^ neurons, which might explain that the mutants develop the clasping phenotype, characteristic for PD-like neurodegeneration and other neurological conditions [79, 80].

To directly prove that the above-mentioned neuroprotection is indeed related to miR-290-295, we aimed to rescue the cluster knockout by overexpression of miR-292a-3p, the most abundant member of the cluster in SN^DA^ neurons (**Figure 2Q**). Following *in vitro* validation of SaCas9-gRNA_290_ and SaCas9-gRNA_295_ (**Figure 2K-R** and **Table S2**), they were subcloned to a construct equipped with a double-floxed inverted open reading frame (DIO)-dependent SaCas9 driven by elongation factor 1α (EFS) promoter to generate a dual-gRNA vector (EFS-DIO-SaCas9-U6-gRNA_290_-U6-gRNA_295_) for the cluster knockout. For miR-292a overexpression, its precursor hairpin was subcloned into the 5′-untranslated region (UTR) of the human synapsin promoter (hSyn)-driven tdTomato (hSyn-DIO-miR-292a-tdTomato). Both of these as well as control Cre-dependent constructs were packaged into neuron-specific CAP.B10 rAAV capsids and stereotaxicly injected into the SNpc of adult *Slc6a3^em(Cre)^* mice to achieve cell-type-specific cluster knockout and/or miR-292a overexpression (**Figure 2S** and **Tables S1-S2**). All injections were performed as 1:1 (vol:vol) mixtures of functional or empty viruses as depicted in **Figure 2Q**. Strikingly, knockout of the cluster impaired the ability of mice to descend from a pole (**Figure S2S**). In our experimental setup, we deliberately aimed for asymmetric alterations, expecting their earlier manifestation, which can be further enhanced by apomorphine, a non-selective agonist of DA receptors. These become hypersensitive on the side of the brain with the DA system deficit, thus resulting in apomorphine-induced contraversive rotations, in a direction opposite to the deficit side. Rewardingly, already at 20 wk after transduction, knockout animals revealed pronounced hindlimb clasping and apomorphine-induced rotations, both of which were successfully rescued in the SN^DA^:miR-290-295CKO_SaCas9_-miR-292a group (**Figures 2T-U, S2T** and **Videos S3-S4**). Accordingly, 2 months later, we detected plummeted mRNAs encoding DDC, TH, and DAT (**Figure 2V-X**) in SN^DA^ neurons of SN^DA^:miR-290-295CKO_SaCas9_ mice accompanied by striatal depletion of the latter two proteins (**Figure 2Y-Z**), all of which were successfully rescued by miR-292a, directly validating the functional role of this microRNA *in vivo*.

### miR-292a-3p is cytoprotective and promotes DA biogenesis

After detecting such striking phenotypes, we next focused on delineating the mechanism of miR-292a-3p. For that, we used the miR-290-295 cluster-expressing MN9D mouse DA neuronal cell line [81] to generate stable miR-292a knock-out clones. First, we predicted, validated and subcloned two gRNAs targeting both strands of miR-292a into the HP180.2 plasmid, which was modified from the three gRNA cassette-containing HP180.3 plasmid [82] equipped with a GFP reporter and SpCas9 (**Figure 3A-C** and **Tables S2-S3**). After transfection, individual GFP-positive MN9D clones were collected by FACS and propagated for subsequent knockout efficiency verification both at the DNA and RNA levels (**Figure 3D-E)** with the clones B3 and F4 selected for further analyses. Interestingly, the latter clone revealed deficits in cell attachment and viability (**Figure 3F-G**), which was rescued by the miR-292a-3p mimic (**Figure 3H**), further proving its causative cytoprotective role. In accordance with our *in vivo* data, deletion of this microRNA decreased the expression of TH, DDC and DAT (**Figure 3I-L**), critical for DA biogenesis, while application of miR-292a-3p rescued both *Th* and *Ddc* levels (**Figure 3M-P** and **Table S2**).

**Figure 3.**
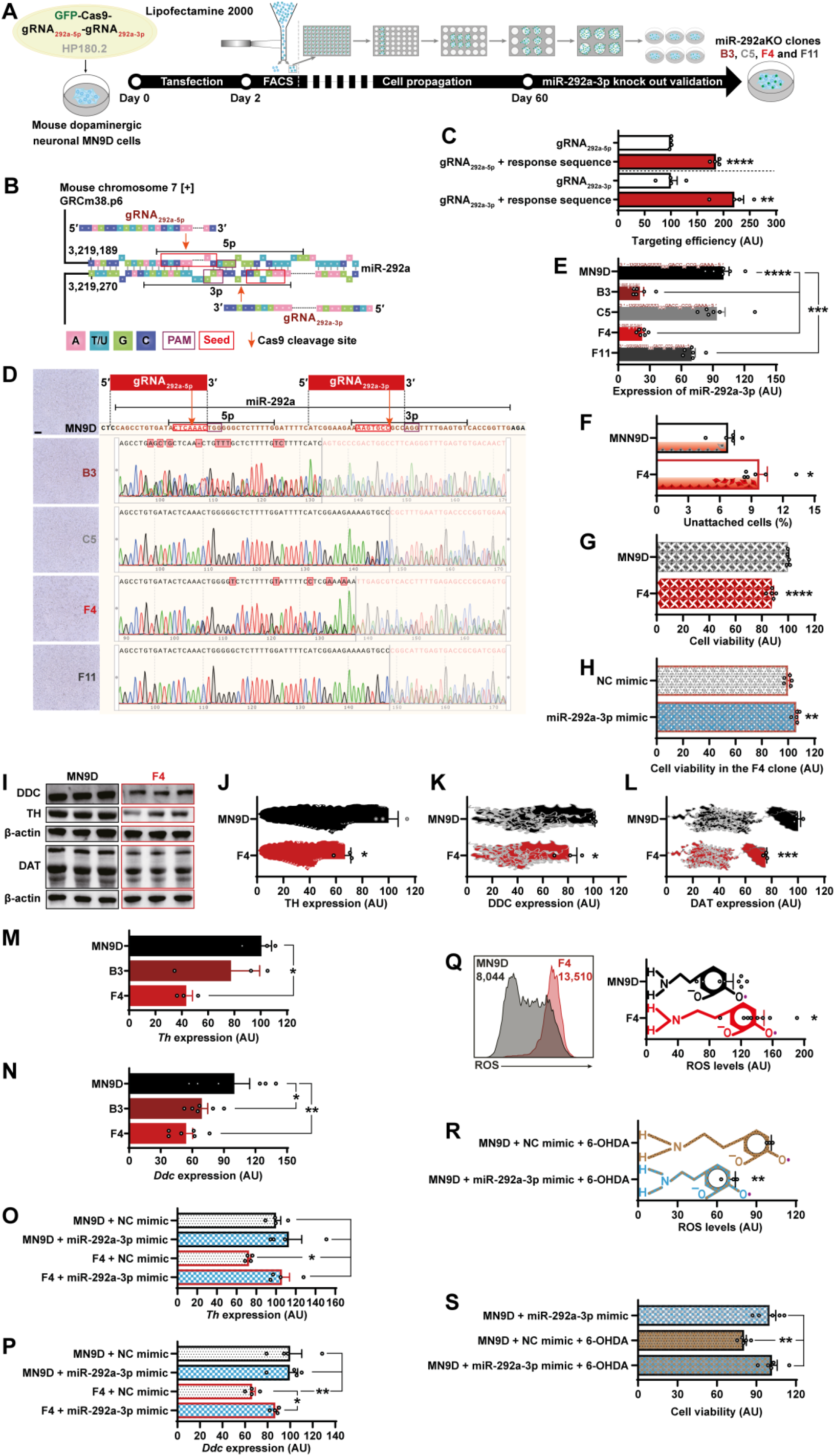
miR-292a-3p is cytoprotective and promotes DA biogenesis. (**A**) Schematic diagram of the procedure to generate miR-292a knock-out clones. (**B-C**) Design (**B**) and on-target efficiency analysis of gRNAs (**C**, n = 4). (**D-F**) Validation of clones by Sanger sequencing (**D**) or qRT-PCR normalized to *U6* (**E**, n = 6) and quantification of attached cells 24 h after replating the F4 clone and intact MN9D cells (**F**, n = 6 and 5, respectively). (**G-H**) Cell viability in the F4 clone (**G**) and its rescue (**H**) after transfection of 30 nM miR-292a-3p mimic (n = 5). (**I-L**) Representative western blot photographs (**I**) and quantification of TH (**J**), DDC (**K**) or DAT (**L**) in the F4 clone and intact MN9D cells normalized to β-actin (n = 3). (**M-P**) *Hprt1*-normalized expression of *Th* (**M**, n = 3) and *Ddc* (**N**, n = 6) in miR-292aKO clones and the rescue of the *B2m*-normalized expression of *Th* (**O**) and *Ddc* (**P**) in the F4 clone after transfection of 30 nM miR-292a-3p mimic (n = 4). (**Q**) Reactive oxygen species (ROS) in the F4 clone and MN9D (n = 8). (**R-S**) Transfection of 30 nM miR-292a-3p mimic reduced ROS production (**R**, n = 3) and increased cell viability (**S**, n = 5) in 6-OHDA exposed MN9D cells. Error bars represent SEM. *, *p* < 0.05; **, *p* < 0.01; ***, *p* < 0.001; ****, *p* < 0.0001 as analyzed by unpaired two-tailed Student’s t-test (**C,F-H,J-L,Q,R**) or one-way ANOVA followed by post-hoc Holm-Šídák’s test (**E,M-P,S**). See the unprocessed western blots and statistical details in the **Source data**.

Due to their electrophysiological and biochemical characteristics, DA neurons are prone to accumulate reactive oxygen species (ROS) with aging or upon degeneration [83]. These reactive products inhibit the mitochondrial Complex I, induce electron leakage, deplete ATP and collapse the inner mitochondrial membrane potential leading to opening of the mitochondrial permeability transition pore (mPTP), which leads to the cytoplasmic release of cytochrome c and caspase- and protein kinase Cδ-mediated apoptosis [84, 85]. Interestingly, miR-292a knockout significantly up-regulated ROS levels in MN9D cells (**Figure 3Q**). Even more potently than DA, its structural analogue 6-hydroxydopamine (6-OHDA), which is widely used in neurodegenerative models, depletes glutathione and impairs the proteasome system via generation of hydrogen peroxide, hydroxyl radicals and quinones [86]. To investigate whether miR-292a-3p can rescue 6-OHDA-induced degeneration, we first validated its ROS-neutralizing function *in vitro* (**Figure 3R**). Strikingly, this mimic also rescued the survival of MN9D cells treated with 6-OHDA (**Figure 3S**). To validate these effects *in vivo*, we stereotaxically injected 6-OHDA to the right striatum of adult *Slc6a3^em(Cre)^* mice with the aim to rescue the asymmetric phenotypes by bilaterally injected miR-292a-overexpressing rAAV (**Figure 4A-B** and **Tables S1-S2**). Despite the absence of pole or clasping phenotypes in 6-OHDA mice (**Figure S3A-B**), our asymmetric paradigm successfully induced contraversive rotations upon treatment with apomorphine. Moreover, similar to the cluster knockout experiments, miR-292a successfully rescued both apomorphine-induced rotations (**Figures 4C**, **S3C** and **Video S5**) and the striatal expression of TH and DAT in these mice (**Figure 4D-G**), further confirming the cytoprotective function of miR-292a-3p in mature DA neurons.

**Figure 4.**
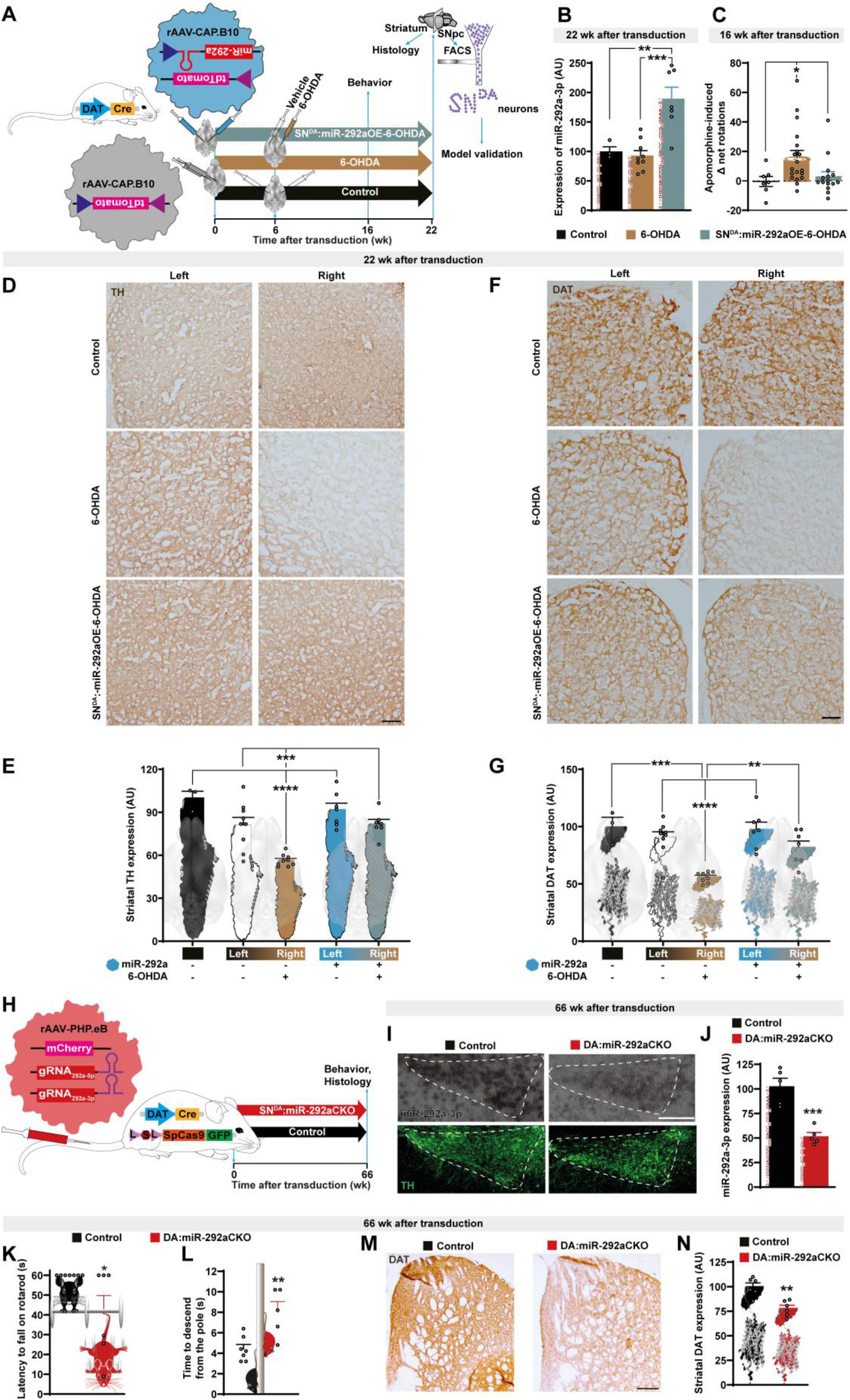
miR-292a protects mature DA neurons *in vivo*. (**A**) Experimental design for assessing the rescue effect of miR-292a overexpression in SN^DA^ neurons following 6-OHDA treatment. rAAV-CAP.B10-miR-292a-tdTomato or rAAV-CAP.B10-tdTomato were stereotaxically injected into both sides of the SNpc across three experimental groups. In the 6-OHDA group, wild type mice (5 females, 2 months; 8 males, 2-5 months) received rAAV-CAP.B10-miR-292a-tdTomato, and *Slc6a3^em(Cre)^* mice (4 females, 3 months; 2 males, 3 months) received rAAV-CAP.B10-tdTomato. In the SN^DA^:miR-292aOE-6-OHDA group, *Slc6a3^em(Cre)^* mice (4 females, 3 months; 10 males, 3-5 months) were injected with rAAV-CAP.B10-miR-292a-tdTomato. In the Control group, wild type mice (5 females, 2 months; 2 males, 5 months) received rAAV-CAP.B10-miR-292a-tdTomato. Six weeks after viral delivery, saline and 6-OHDA were stereotaxically injected into the left and right striatum, respectively, for the 6-OHDA and SN^DA^:miR-292aOE-6-OHDA groups, whereas the Control group received saline bilaterally. (**B**) Expression of miR-292a-3p in SN^DA^ neurons sorted from 6-OHDA, SN^DA^:miR-292aOE-6-OHDA and Control groups (n = 9, 7 and 3, respectively) 22 wk after transduction. (**C**) Apomorphine-induced Δ net rotations in 6-OHDA, SN^DA^:miR-292aOE-6-OHDA and Control mice (n = 19, 15 and 7, respectively) 16 wk after transduction. Δ net rotations = number of full 360° contraversive - number of full 360° ipsiversive rotations. (**D-G**) Representative microphotographs (**D,F**) and quantification (**E,G**) of striatal immunostaining for TH (**D,E**) and DAT (**F,G**) in 6-OHDA, SN^DA^:miR-292aOE-6-OHDA and Control mice (n = 9, 7 and 3, respectively) 22 wk after transduction. (**H**) Experimental design to delete miR-292a in mature DA neurons. For the DA:miR-292aKO group, rAAV-PHP.eB-gRNA_292a-5/3p_ was injected via tail vein into 3-4-month-old DAT^Cre^:SpCas9-GFP mice (5 females and 4 males). For the Control group, rAAV-PHP.eB-empty was injected via tail vein into 3-4-month-old wilde type mice (6 females and 4 males). (**I-J**) Representative microphotographs (**I**) and quantification (**J**) of miR-292a-3p *in situ* hybridization in TH-positive neurons within the SNpc (n = 5) to validate the knockout efficiency in DA:miR-292aCKO mice. (**K-L**) Performance of DA:miR-292aCKO and Control mice on the rotarod (**K**, n = 6 and 7, respectively) and pole assays (**L**, n = 5 and 7, respectively) 66 wk after transduction. (**M-N**) Representative microphotographs (**M**) and quantification (**N**) of striata immunostained for DAT (n = 6). Error bars represent SEM. *, *p* < 0.05; **, *p* < 0.01; ***, *p* < 0.001; ****, *p* < 0.0001 as analyzed by one-way ANOVA followed by post-hoc Holm-Šídák’s test (**B,E,G**), Kruskal-Wallis test with Dunn’s multiple comparisons test (**C**), unpaired two-tailed Student’s t-test (**J,L,N**), or Log-Rank test (**K**). See the details about ages, sexes, FACS gating strategy and statistical methods in the **Source data**. Scale bar: 200 μm.

### Loss of miR-292a in mature DA neurons leads to aging-associated locomotor and striatal DAT deficits

To complement these gain-of-function data, we intravenously injected adult Dat^Cre^:SpCas9-GFP mice with gRNA_292a-5/3p_-equipped PHP.eB-rAAV vector which is capable to cross the blood-brain barrier and knock-out this microRNA in mature DA neurons (**Figures 4H-J**, **S3D** and **Tables S1-S3**). While DA neuron-restricted deletion of only one microRNA did not induce clasping behavior up to 66 wk after transduction (**Figure S3E**), we aimed to use the pole test, which, due to the functional constraints, is particularly applicable when no clasping is observed [87]. Strikingly, animals with miR-292a deficiency revealed age-dependent deficits in both pole climbing and performance on the originally developed modified constant speed rotarod, one of the most sensitive behavioral paradigms for detecting neurodegenerative phenotypes [61] (**Figures 4K-L** and **S3F**). These late onset locomotor deficits in DA:miR-292aCKO mice were associated with the decline in striatal expression of DAT (**Figure 4M-N**), but neither TH nor DDC (**Figure S3G-J**), nor the loss of SN^DA^ neurons (**Figure. S3K**), nor the projection density in the *Substantia Nigra pars reticulata* (**Figure S3L-M**) 15 months after transduction. To determine whether DA neurons from the VTA contributed to the observed striatal decline in DAT immunoreactivity, we further quantified the number of these cells and their projections to the ventral striatum, discriminating this region from the dorsal striatum, where the SN^DA^ neuronal terminals predominantly reside. Notably, neither the VTA^DA^ neuronal counts nor their projection density revealed significant alterations in DA:miR-292aCKO mice (**Figure S3N-O**). These data suggest that the VTA^DA^ neurons did not contribute to or confound the observed phenotypes. Altogether, our results delineate a protective function of miR-292a in mature DA neurons, while suggesting the dosage-dependent effect when DA neuron-restricted deletion of multiple microRNAs sharing the same seed region may cause an earlier onset of a more pronounced phenotype compared to an isolated deletion of miR-292a.

### *Pten*, a direct target of miR-292a-3p, is drastically up-regulated in SN^DA^ neurons of adult SN^DA^:miR-290-295CKO mice

To prove the above-mentioned hypothesis that the common seed region shared by the members of the cluster may target genes capable of driving the above-mentioned phenotypes, we sought to identify such common direct targets. First, we performed the Smart-seq2 analysis on FACS-isolated SN^DA^ neurons from adult SN^DA^:miR-290-295CKO mice and used microT-CDS v. 5.0 [88] to predict which of the top-elevated genes are targeted by all six major microRNAs in the cluster (**Figure 5A-C** and **Source data**). Strikingly, out of the top 30 up-regulated genes, phosphatase and tensin homolog (*Pten*) was the only putative target of all these microRNAs. This gene encodes for PTEN, a major antagonist of the phosphatidylinositol-4,5-bisphosphate 3-kinase (PI3K)-AKT pathway, which is upstream of the mechanistic target of rapamycin kinase (mTOR) pathway, both of which were inhibited in SN^DA^ neurons of SN^DA^:miR-290-295CKO mice (**Figure 5D**). mTOR complex 1 (mTORC1) stimulates cell growth and protein synthesis by activating S6 ribosomal subunit and eukaryotic translation initiation factors B and E (eIF4B and eIF4E, respectively). Accordingly, *Eif4e* encoding this key factor was the most down-regulated gene in SN^DA^ neurons of SN^DA^:miR-290-295CKO mice (**Figure 5A**) pointing together with la ribonucleoprotein 4 (*Larp4*) and reciprocally up-regulated CCR4-Not transcription complex subunit 7 (*Cnot7*) and tuberous sclerosis complex 1 (*Tsc1*) to a drastic inhibition of the PI3K-AKT-mTORC1 pathways, impairment of protein synthesis and, conversely, activation of the downstream FoxO1 pathway (**Figure 5D**), which is known to directly inhibit TH expression [89]. Indeed, expression of both TH and DAT can be stimulated by the survival-promoting PI3K-AKT pathway [90, 91], which in turn is strongly antagonized by PTEN. Using the dual luciferase reporter system, we have proven all three predicted binding sites on the 3′-UTR of *Pten* as direct targets of miR-292a-3p, the most abundant SN^DA^ neuron-expressed microRNA from this cluster (**Figure 5E-H**).

**Figure 5.**
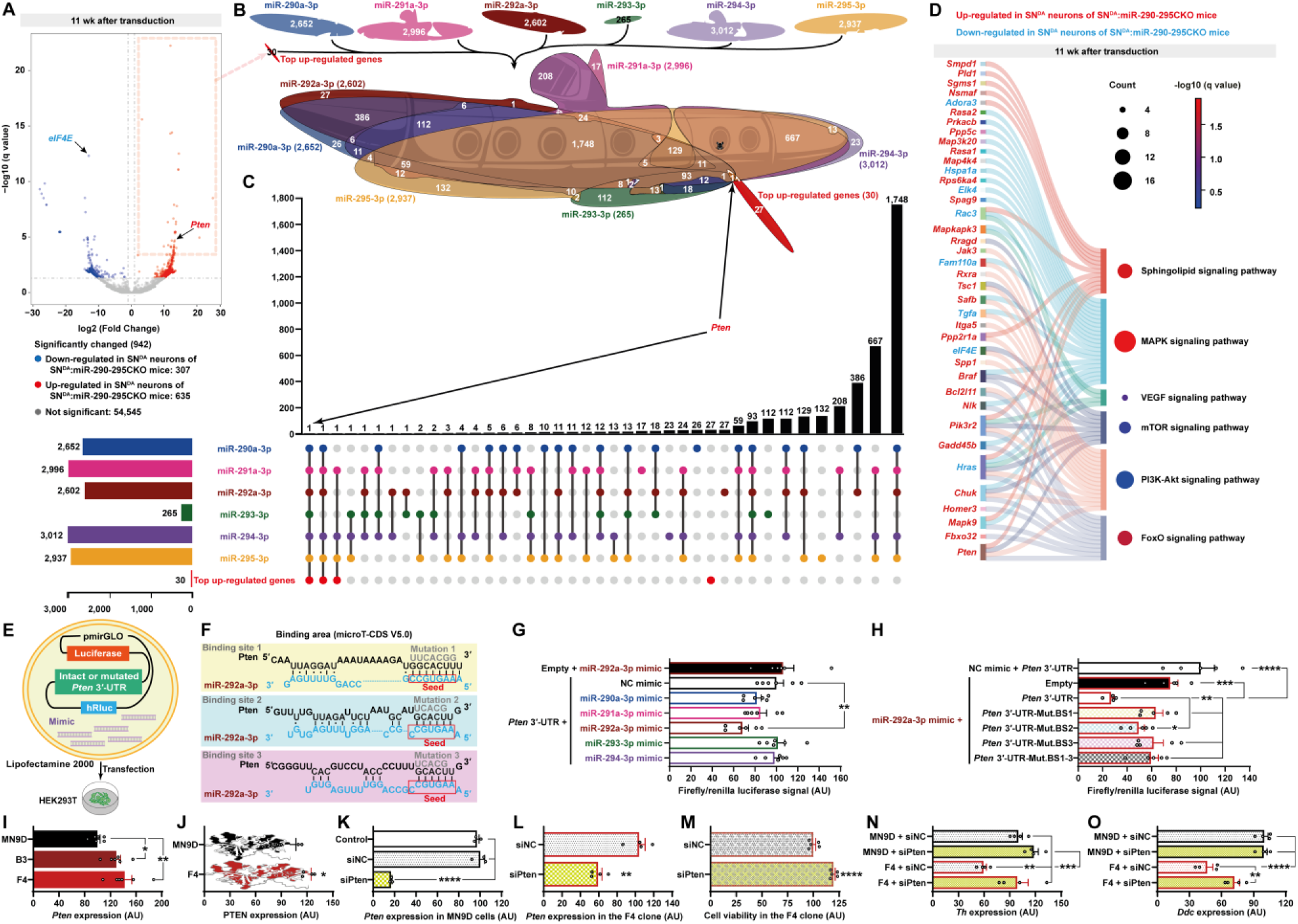
*Pten*, a direct target of miR-292a-3p, is drastically up-regulated in SN^DA^ neurons of adult SN^DA^:miR-290-295CKO mice. (**A**) Volcano plot of Smart-seq2 demonstrating significantly changed genes in FACS-isolated mature SN^DA^ neurons of SN^DA^:miR-290-295CKO mice (1 female aged 13-months, 1 female aged 9-months and 1 male aged 15-month) compared to Controls (1 male aged 15-months and 2 males aged 13-months) 11 wk after transduction. (**B-C**) Euler diagram approximation (**B**) and UpSet plot (**C**) identifying phosphatase and tensin homolog (*Pten*) as the only candidate target of the depicted cluster microRNAs within the genes mostly upregulated in mature SN^DA^ neurons of SN^DA^:miR-290-295CKO mice. (**D**) Kyoto Encyclopedia of Genes and Genomes-assisted analysis of the most affected signaling pathways in mature miR-290-295 cluster-depleted SN^DA^ neurons. (**E-G**) Experimental design of the dual-luciferase target validation assay (**E**), putative *Pten* binding sites and mutated sequences used in this assay (**F**) and quantification (**G**) of *Pten* validation as a target (n = 6). (**H**) Further validation using vectors with mutated binding sites (n = 5 except the miR-292a-3p mimic-Pten 3′-UTR group, which had n = 4). (**I**) Expression of *Pten* in miR-292aKO clones (n = 6) normalized to *Hprt1*. (**J**) Expression of PTEN in F4 clone (n = 4). (**K-L**) Validation of the *Pten*-targeting siRNA efficiency in intact (**K**, n = 3) and miR-292a-deficient MN9D cells (**L**, n = 4) normalized to *Hprt1* and *B2m*, respectively. (**M-O**) Cell viability (**M**, n = 5), *B2m*-normalized expression of *Th* (**N**) and *Ddc* (**O**) in the F4 miR-292aKO clone (n = 4) after transfection with 800 nM siRNA targeting *Pten*. Error bars represent SEM. *, *p* < 0.05; **, *p* < 0.01; ***, *p* < 0.001; ****, *p* < 0.0001 as analyzed by one-way ANOVA followed by post-hoc Holm-Šídák’s test (**G-I,K,M-O**), or unpaired two-tailed Student’s t-test (**J,L**). See the details about FACS gating strategy, quality control data for Smart-seq2 and statistical methods in the **Source data**.

### miR-292a-3p targets *Pten* to maintain cell viability and DA biogenesis

To validate the function and mechanism of the miR-292a-PTEN axis in MN9D cells, we sought to knock-down *Pten* in miR-292a-deficient clones. While, similar to the above *in vivo* data, PTEN was up-regulated in the mutant cells (**Figure 5I-J**), application of *Pten*-targeting siRNA successfully inhibited it in both intact and mutant MN9D cells (**Figure 5K-L**). Strikingly, this siRNA restored both survival and expression of *Th* and *Ddc* in miR-292a-deficient cells, with the latter gene being significantly up-regulated in comparison to the non-targeting control siRNA, but also revealing a downward tendency vs. the non-mutated MN9D cells, which, however, did not reach a statistical significance (**Figure 5M-O**). These data validate the cytoprotective mechanism of miR-292a via inhibition of its direct target PTEN.

### The miR-290-295 cluster maintains protein synthesis in mature SN^DA^ neurons

Previously, we have demonstrated an enormous beneficial effect of PTEN inhibition in mature midbrain DA neurons, leading to increased phosphorylation of AKT and S6 ribosomal proteins in this region, which reflects a strong activation of the neuroprotective PI3K-AKT-mTORC1-protein synthesis pathways. Importantly, knock-out of PTEN in DA neurons subsequently promotes DA synthesis and reuptake, indicated by increased expression of TH and DAT, while counteracting neurodegeneration [91]. As discussed above, SN^DA^ neurons from SN^DA^:miR-290-295CKO mice revealed a drastic up-regulation of *Pten* and, consistently, an alteration of the PI3K-AKT-mTORC1-protein synthesis signaling pathways (**Figure 5A,D**).

Next, we sought to directly prove that among various pathways and processes regulated by the PTEN-PI3K antagonism, protein synthesis is essential for maintained DA biogenesis. For that, we used an adult-onset model with a knock-out of RNApol I-specific transcription initiation factor Rrn3 in mature DA neurons, which alters their nucleoli, ribosomal RNA synthesis and hence translation. We sought to test whether this protein synthesis deficit abolishes the previously reported PTEN loss-mediated up-regulation of TH, the rate-limiting enzyme in DA biosynthesis [91]. Indeed, we failed to detect an enhanced TH expression in Dat^Cre-ERT2^:Rrn3^fl/fl^Pten^fl/fl^ compared to Dat^Cre-ERT2^:Rrn3^fl/fl^ mice (**Figure 6A-C**), suggesting that the defect in protein synthesis profoundly disrupts DA neuronal metabolism, such as that even activation of the PI3K pathway and its downstream targets is unable to rescue it.

**Figure 6.**
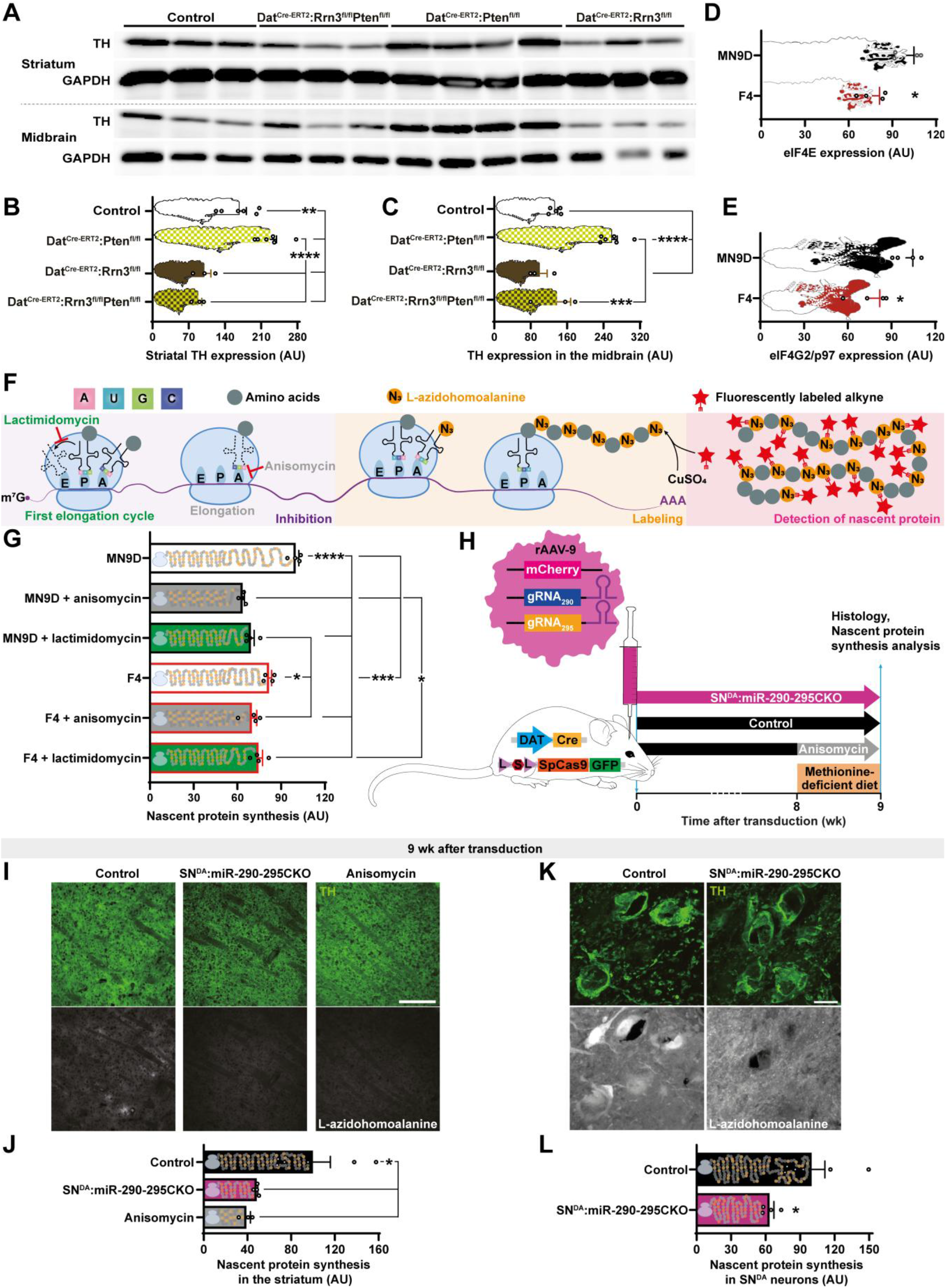
miR-290-295 maintains protein synthesis *in vivo*. (**A-C**) Representative western blot photographs (**A**) and quantification of TH expression in the striatum (**B**, n = 6, 3, 3 and 6, respectively) or the midbrain (**C**, n = 6, 3, 3 and 5, respectively) of Dat^Cre-ERT2^:Pten^fl/fl^, Dat^Cre-ERT2^:Rrn3^fl/fl^, Dat^Cre-ERT2^:Rrn3^fl/fl^Pten^fl/fl^ and Control mice normalized to GAPDH. (**D-E**) Expression of eukaryotic translation initiation factors E (eIF4E, **D**) and eIF4G2/p97 (**E**) in the F4 clone (n = 4). (**F**) Scheme illustrating the mechanism of protein inhibitor action and the application of L-azidohomoalanine for detection of nascent proteins. (**G**) Nascent protein synthesis in the F4 clone and MN9D (n = 4). (**H**) Experimental design of the nascent protein synthesis assay by i.p. injection of click chemistry-compatible methionine agonist L-azidohomoalanine into methionine-deprived SN^DA^:miR-290-295CKO mice with or without prior injection of protein synthesis inhibitor anisomycin. rAAV-9-gRNA_290/295_ was stereotaxically injected into SNpc of 7-10-month-old DAT^Cre^:SpCas9-GFP mice (4 females and 1 male). For the Control group, empty rAAV was injected into SNpc of 4-15-month-old DAT^Cre^:SpCas9-GFP mice (3 females and 4 males). For the Anisomycin group, this inhibitor was injected into 6 wild type female mice aged 4-15 months. (**I-L**) Representative microphotographs of newly synthesized proteins (**I,K**) and quantification (**J,L**) in the striatum (**I,J**) or SN^DA^ neurons (**K,L**) of SN^DA^:miR-290-295CKO, Control and Anisomycin negative control mice (n = 4, 6 and 3, respectively). Error bars represent SEM. *, *p* < 0.05; **, *p* < 0.01; ***, *p* < 0.001; ****, *p* < 0.0001 as assessed by one-way ANOVA with post-hoc Holm-Šídák’s test (**B,C,G,J**) or unpaired two-tailed Student’s t-test (**D,E,L**). Scale bars in µm: 200 (**I**), 20 (**K**). See unprocessed western blots, details about ages, sexes and statistical methods in the **Source data**.

In agreement with our *in vivo* data, miR-292a-deficient MN9D cells revealed a down-regulation of both translation initiation factors eIF4E and eIF4G2/p97 (**Figure 6D-E**), further indicating the impairment of this stage of protein synthesis. eIF4E mediates binding of the eIF4F complex to the mRNA 5′ cap structure, thus governing the translation initiation, the rate-limiting step of protein synthesis. To directly demonstrate the causal role of miR-292a in protein synthesis and, more specifically, translation initiation, we employed the Fluorescent Noncanonical Amino Acid Tagging (FUNCAT) approach [92]. Implementing a click chemistry-compatible methionine analogue L-azidohomoalanine to label the newly synthesized proteins in MN9D cells, we then treated them with two translation inhibitors targeting different stages of translation (**Figure 6F**). While an elongation inhibitor anisomycin interferes with the late stages of translation involving peptide bond formation and peptidyl transferase activity on the 60S ribosomal subunit, lactimidomycin targets the E-site of the same subunit to block the early translocation step of the first elongation cycle. This allowed us to differentiate, whether the early or late translation stage could still be impaired by these inhibitors beyond the extent caused by the knockout of the PTEN-targeting microRNA. Strikingly, while the miR-292a-deficient F4 clone revealed diminished nascent protein synthesis, anisomycin, but not lactimidomycin could further impair translation, in contrast to the untargeted MN9D cells, where both inhibitors were effective to suppress protein synthesis (**Figure 6G**). These data directly indicate the causative role of miR-292a in preservation of protein synthesis via promoting the early translation stages, as demonstrated by inability of lactimidomycin to further suppress it, as well as based on our *in vitro* and *in vivo* analyses of translation initiation factor expression.

Finally, to confirm the impairment of nascent protein synthesis upon loss of miR-290-295 *in vivo*, we applied the FUNCAT approach to SN^DA^:miR-290-295CKO or Control mice fed with methionine-deficient diet [93] with or without application of anisomycin as a negative control (**Figure 6H**). In agreement with our previous data, as early as 2 months after transduction, SN^DA^:miR-290-295CKO mice revealed reduced DAT levels in the striatum and the medial forebrain bundle, but not the *Substantia Nigra* (**Figure S4**). These results point to an alteration in DA biogenesis within the projections of SN^DA^ neurons. Strikingly, upon miR-290-295 knock-out, we detected a drastic down-regulation of protein synthesis in both SN^DA^ neurons (colocalized with TH) and striatum to the same level as in the negative control animals treated with anisomycin (**Figure 6I-L**). These and previous data suggest that a sustained damage to the PI3K-AKT-mTORC1-protein synthesis pathways gradually leads to deficits in DA biosynthesis and SN^DA^ fitness, as well as impairment of neuronal projections, already within 9 wk after transduction.

Altogether, we’ve identified the physiological role and the mechanism of the most DA neuron-enriched microRNAs in SNpc. Keeping PTEN in check by the miR-290-295 cluster assures adequate protein synthesis levels which are essential for DA biogenesis and neuroprotection in the adulthood. Conversely, knockout of the miR-290-295 cluster in mature SN^DA^ neurons leads to down-regulation of the PI3K-AKT-mTORC1-protein synthesis pathways, compromising DA biogenesis and contributing to molecular and behavioral hallmarks of neurodegeneration.

## DISCUSSION

For the first time, we report that adult SN^DA^ neurons express the stemness-specific miR-290-295 cluster, which reveal an age-related decline. Intriguingly, these are also the most SN^DA^ neuron-enriched microRNAs in this region, playing a critical role in DA biogenesis and SN^DA^ neuronal fitness. Indeed, the global knock out of miR-290-295 leads to an allelic dosage-dependent decline of SN^DA^ neuronal numbers in adult mice, in agreement with our *in vitro* data that both cell survival and DA biogenesis are dependent on the expression of the most abundant cluster member miR-292a-3p, which antagonizes its direct target PTEN. Deletion of the cluster or miR-292a from DA neurons of adult mice induces both molecular and behavioral hallmarks of neurodegeneration, including early onset down-regulation of key players involved in the DA biogenesis within the SN^DA^ neurons and their projections, but not the loss of their cell bodies. Derepression of PTEN upon miR-290-295 deletion from SN^DA^ neurons leads to a marked suppression of mTORC1-controlled eIF4E, a key factor for cap-dependent translation initiation, causing alteration of protein synthesis, which, in turn, is essential for DA biogenesis.

Depending on the quantification method, SNpc comprises 47 to 74% SN^DA^ neurons, which are supported by other neuronal (up to 43%) and non-neuronal cell populations [94], such as interneurons, astrocytes, microglia, oligodendrocytes, endothelial, mural and ependymal cells [73, 95–98]. In this work, we identified microRNAs highly enriched or exclusive to SN^DA^ neurons. For that, we designed a screening approach on the mice with DA neuron-specific knock-out of Dicer or their control littermates. To selectively label SN^DA^ neurons, we utilized fluorescent retrobeads injected into the striatum. The SNpc tissue for microRNA profiling was dissected encircling the whole group of retrobead-positive neuronal bodies in each section, thus including both SN^DA^ neurons and the neighboring cells. As such, for microRNAs not enriched in SN^DA^ neurons, the higher their abundance in non-DA cells and the higher the number of such cells within the dissected SNpc tissue, the less sensitive TaqMan qRT-PCR would be to DA neuron-restricted Dicer knock-out, due to the masking of microRNA decline by the surrounding cells with intact Dicer. Indeed, our Dicer loss-of-function model demonstrated not only the abundance, but also a remarkable DA neuron-enrichment of all cluster microRNAs, due to the dependence of their maturation on Dicer expression [99, 100].

Notably, the rationale for using different time points after Dicer knockout was related to the dynamics of previously observed molecular and behavioral alterations [61]. Despite a significant decline 2 weeks after tamoxifen injections, *Dicer1* mRNA was not completely degraded in DA neurons of Dat^Cre-ERT2^:Dicer^fl/fl^ animals, which suggests a retention of a sufficient pool of Dicer protein to maintain, at least partially, its critical function in microRNA biogenesis. The following depletion of striatal dopamine content, the very first molecular hallmark of neurodegeneration in these mice, was revealed as early as on Week 4, which was chosen as the earliest time point for the enrichment experiment in this study. This ensured the down-regulation of microRNAs, especially those ones critical for DA neuron function. Still, we could not exclude the possibility that some traces of *Dicer1* mRNA and Dicer protein still retained in these neurons at this time point. As such, we have added the Week 6 time point, when the striatal tyrosine hydroxylase and the number of DA cells themselves started to decline both in the *Substantia Nigra* and the VTA of Dat^Cre-ERT2^:Dicer^fl/fl^ mice [61]. We did not risk to wait longer, since the fulminant course of neurodegeneration in these animals results in devastation of the *Substantia Nigra* and immobility as early as by Week 10. As such, the strategy of dissecting and profiling this region could yield confusing or uninterpretable results due to the loss of the very cell population being studied.

To our knowledge, such loss-of-function approach has never been used before to detect the most enriched transcripts in a defined cellular population. It opens a possibility for other research groups to use similar unconventional logic to address other biological problems. Enrichment within a target cell population may mean evolutionary conservation of epigenetic disinhibition locally, with simultaneous silencing in the surrounding tissue, which may point to a functional relevance of that gene for this cell. Notably, the strongest enrichment of a microRNA in this cell population within a defined region doesn’t mean the highest expression of this microRNA in this cell. Functionally, microRNAs representing the largest share of the whole microRNA pool play the major role in a specific cell lineage, exemplified by miR-290-295 constituting 2/3 of all microRNAs in embryonic stem cells [8–13]. This is opposed to functional insignificance of the minor microRNAs [101] which fail to outcompete not only the major ones to occupy AGO2, but also messenger RNA targets and pseudo-targets to exert meaningful silencing [102]. Mechanistically, largest microRNA fractions occupy the limited number of argonaute 2 (AGO2) within the RNA-induced silencing complex (RISC), thus exhausting the pool of these proteins for other minor microRNAs [103]. This translates to only few convincing reports about physiological functions of minor microRNAs expressed locally or introduced via extracellular vesicles from other cells [104–107]. As such, identifying the most cell population-enriched microRNAs is just the first step towards functional and mechanistical assessment. The next logical step requires identification of expression levels within the defined cell population, followed by functional characterization targeting the most abundant microRNAs. While the stem-loop primer-based qRT-PCR is the golden standard to quantify the relative expression of mature microRNAs, a direct comparison of abundances between different microRNA species is unreliable when assessed with intercalating dyes, such as SYBR Green. This is because the dynamics of quantification curves depends not only on the transcript abundance, but also on the GC content, amplicon length and sequence arrangement, precluding direct quantitative comparisons between different microRNA within the same sample. In contrast, TaqMan hydrolysis probe chemistry introduces an indispensable layer of sequence specificity while ensuring a 1:1 stoichiometric fluorophore release per amplicon. Since fluorescence is emitted exclusively upon target-specific probe hybridization and cleavage, this approach effectively eliminates background artifacts arising from non-specific amplification or primer-dimers, thereby markedly improving quantification accuracy across distinct targets. Accordingly, in our study, after identification of miR-290-295 as the most enriched microRNA in SN^DA^ neurons within SNpc, we next determined the dominant species among these, miR-292a-3p. Indeed, as determined by TaqMan qRT-PCR, it was similarly abundant in SN^DA^ neurons as the previously reported DA neuron-expressed miR-133b and miR-124. This placed this microRNA among the most expressed microRNAs within these cells, thus identifying it as our primary candidate for further functional studies.

Importantly, in addition to Dicer loss-of-function model, we confirmed the abundance and/or enrichment of the cluster microRNAs within the mature SN^DA^ neurons by four complementary techniques: FACS-sorting of retrobead- or DAT-positive neurons, laser capture-microdissection and *in situ* hybridization. While the sensitivity of these methods vary due to the insufficiency of the total RNA amount from the limited number of collected cells or specificity of the probes, such multimodal confirmation tremendously strengthens our conclusions. Notably, pooling several mice per replicate allowed us to collect sufficient number of cells for FACS-assisted techniques. Interestingly, in addition to the cluster microRNAs which are highly enriched or exclusive to SN^DA^ neurons in this region, we detected several other enriched microRNAs, such as miR-504-5p, miR-139-3p, miR-302 and miR-409-5p. However, only miR-290-295 and miR-504-5p qualified our 3-fold expression loss cutoff at both 4 and 6 weeks after Dicer loss. According to our unpublished data, miR-504-5p is also the most down-regulated microRNA in SN^DA^ neurons of patients with idiopathic PD, while mature SN^DA^ neuron-specific knockout of miR-504 leads to locomotor deficits, making this microRNA a promising candidate for further analysis. Abundant in various brain regions, such as cortex or hippocampus, miR-504-5p targets D1 receptor (*DRD1*) [108], *p39* [109], reduces dendritic spine density via targeting SH3 and multiple ankyrin repeat domains 3 (*SHANK3*) [110] and is considered for therapeutic strategies to treat depression-like behaviors [111] or Alzheimer’s disease [109].

The initial molecular hallmarks of neurodegeneration usually manifest in the neuronal terminals [112–115], only later leading to DA neuronal loss [112]. Accordingly, in contrast to the global cluster knockout demonstrating a drastic decline in SN^DA^ neuronal numbers, mature DA neuron-specific knock-out of the cluster microRNAs revealed the hallmarks of DA neuronal degeneration only in the striatum, except for the depletion of mRNAs critical for the DA biogenesis in SN^DA^ neurons. Interestingly, i). the earlier onset—just two months after transduction of SN^DA^:miR-290-295CKO animals —and ii). higher severity (down-regulation of both TH, DDC and DAT) of this molecular phenotype in SN^DA^:miR-290-295CKO and SN^DA^:miR-290-295CKO_SaCas9_ vs. DA:miR-292aCKO mice suggest the dosage-dependent effect upon depletion of multiple microRNAs sharing the same seed region vs. only miR-292a. Throughout this work, we used three independent Cre transgenic models to achieve developmentally uncoupled dopaminergic recombination: Dat^Cre-ERT2^, as well as two Dat^Cre^ lines, B6.SJL-*Slc6a3^tm1.1(cre)Bkmn^*/J and *Slc6a3^em(Cre)^*. The widening of mechanistic insights and the reproducibility of phenotypes across these diverse genetic or viral substantially strengthens the rigor of our conclusions, demonstrating that the observed phenotypes reflect genuine biological consequences rather than model-specific effects. Similarly, we also deliberately used multiple loss-of-function genetic models with various kinds of delivery routes, capsids, CRISPR-Cas9 system types, anatomic distribution of transduced cells to demonstrate that, despite variations of onsets or extents, the resulting phenotypes are highly reproducible and robust. Indeed, the key conclusions of this study were strengthened by the following five knockout models: i). developmentally uncoupled tamoxifen-dependent deletion of all Dicer-dependent microRNAs in mature dopamine neurons; ii). global knockout of the whole cluster starting from the embryonic development stages; iii). developmentally uncoupled SpCas9-dependent deletion of the whole cluster specifically in adult *Substantia Nigra* dopamine neurons; iv). developmentally uncoupled SaCas9-dependent deletion of the whole cluster specifically in adult *Substantia Nigra* dopamine neurons with and without the flip-excision (FLEX)-dependent rescue approach and v). developmentally uncoupled SpCas9-dependent deletion of the miR-292a in mature dopamine neurons.

Importantly, these loss-of-function models were complemented by developmentally uncoupled overexpression of miR-292a specifically in adult *Substantia Nigra pars compacta* dopamine neurons. This gain-of-function model allowed us to use the rescue approach involving Cre-dependent miR-292a-3p up-regulation to demonstrate the specificity of the knockout, as well as to further validate the protective role of this microRNA in DA biogenesis and physiology of these neurons. Even more striking is the finding that miR-292a overexpression rescues both the behavioral and molecular hallmarks of neurodegeneration in the 6-OHDA model. These rescue effects are mechanistically consistent with the established anti-apoptotic function of the miR-290-295 cluster. Indeed, these microRNAs suppress genotoxic-stress-induced apoptosis by directly targeting caspase-2 (*Casp2*) and etoposide-induced protein 2.4 homolog (*Ei24*). Thereby miR-290-295 constrains p53-dependent caspase-3 activation [11] in response to 6-OHDA-induced ROS accumulation, protein oxidation, DNA damage and activation of the p53-BCL2 apoptosis regulator-binding component 3 (BBC3, or PUMA) axis together with unfolded protein response (UPR)-driven BCL-2 homologous antagonist/killer 1 (BAK1) oligomerization and cytochrome c release [116–118].

Among the top-elevated genes in SN^DA^ neurons of SN^DA^:miR-290-295CKO mice, *Pten* was predicted as the only strong target of microRNAs in this cluster, which was also confirmed experimentally for miR-292a-3p. We and others previously demonstrated PTEN as a key suppressor of the neuroprotective PI3K-AKT-mTORC1-protein synthesis pathways in mature DA neurons [91, 119–121]. Previously, we demonstrated that deletion of PTEN in mature DA neurons increases their fitness and protects from neurodegeneration via drastic enhancement of the PI3K-AKT-mTORC1-protein synthesis pathways, as evidenced by increased phosphorylation of AKT and S6 ribosomal protein [91]. Accordingly, the deletion of PTEN protects DA neurons in mitochondrial transcription factor A (Tfam)-deficient (*DAT^Cre^:Tfam^fl/fl^*) MitoPark mice which exhibit parkinsonism-like phenotype [121]. Indeed, PTEN-depleted mice reveal elevated DA content, hypertrophy, increased number and arborization of DA neurons, as well as elevated levels of TH and DAT, the critical proteins for DA biogenesis [91, 120, 121]. Mechanistically, we previously demonstrated that activation of PI3K-AKT-mTORC1-protein synthesis pathways in response to PTEN deletion results in up-regulation of *Foxa2*, *Pitx3*, *En1*, *Nr4a2* and *Lmx1b* encoding proteins critical both for the development and maintenance of DA neurons [50, 51, 53, 54, 91, 122].

The PTEN-PI3K-AKT-mTOR axis has emerged as a central hub for dopaminergic neuron protection across multiple injury paradigms. Beyond genetic deletion of PTEN, sustained pharmacological or viral AKT activation, consistently safeguards both somata and axons. For instance, the small-molecule AKT agonist SC79 effectively prevents oxidative-stress-induced apoptosis and necrosis in DA neurons [123], confirming that direct AKT activation is sufficient to maintain neuronal viability under toxic stress. Similarly, AAV-mediated delivery of constitutively active myristoylated AKT (Myr-AKT) protects nigral cell bodies and axons in 6-OHDA-lesioned mice [124, 125]. Notably, even after axonal degeneration, delayed expression of Myr-AKT or constitutively active Ras homolog enriched in brain (RHEB), an upstream mTOR activator, can drive de novo axonal regrowth from the SNpc to the striatum, enabling robust and long-lasting regeneration of the adult nigrostriatal pathway [126].

MicroRNA-based strategies converge on the same PTEN-PI3K-AKT-mTOR pathways, offering a complementary therapeutic avenue. In a stroke model, systemic delivery of exosomes enriched with the miR-17-92 cluster markedly enhances axonal and dendritic remodeling, synaptic plasticity, oligodendrogenesis and functional recovery via PTEN suppression and subsequent activation of AKT, mTOR, and inhibitory phosphorylation of glycogen synthase kinase-3 (GSK-3β) [127]. Interestingly, delivery of bone marrow mesenchymal stem cell-derived small extracellular vesicles containing the mixture of miR-292, miR-92a and miR-182 (but not any single microRNA thereof) promote survival and regeneration of injured retinal ganglion cells by down-regulating PTEN expression [128], highlighting a broader neuroprotective role for PTEN-targeting microRNAs across neuronal types. Altogether, evidences from genetic, pharmacological, and microRNAs-based interventions consistently demonstrate that strategies preserving axonal integrity or restoring dopaminergic connectivity hold promise as potential therapeutic approaches. Given that axonal degeneration often precedes and exceeds neuronal body loss in many neurodegenerative conditions, stabilizing vulnerable axons or re-establishing their projections represents a mechanistically grounded and clinically meaningful avenue for neuroprotection [129].

In agreement with these *in vivo* data, supplementation of miR-292a-3p or down-regulation of its direct target PTEN rescued the viability and factors critical for dopamine biogenesis in miR-292aKO clones. Strikingly, *Eif4e*, the most down-regulated gene in SN^DA^ neurons of SN^DA^:miR-290-295CKO mice, encodes for the key m^7^G cap-binding protein and recruits ribosomes for the critical regulatory step in translation initiation [130, 131]. eIF4E is not only essential for cap-dependent translation initiation but also regulates multiple mRNA-level processes, including RNA capping, splicing, polyadenylation, and selective nuclear export of transcripts containing the eIF4E-sensitive element via the leucine-rich pentatricopeptide repeat-containing protein (LRPPRC)-chromosomal maintenance 1/exportin-1 (CRM1/XPO1) pathway. Furthermore, cytoplasmic interactions of eIF4E with eIF4G, cytoplasmic fragile X messenger ribonucleoprotein 1 (FMR1)-interacting protein 1 (CYFIP1) and eIF4E binding protein 1 (4E-BP1) not only stimulate translation, but also stabilize transcripts [132], hence its cluster knockout-dependent down-regulation might contribute to the observed decline in mRNA levels of both *Th*, *Dat*, *DDC* and *Eif4e* itself. Furthermore, consistent with our data, PI3K-AKT pathways have been reported to indirectly stabilize *eIF4E* transcript in a ELAV-like RNA-binding protein 1 (ELAVL1, or HuR)-dependent manner. Indeed, AKT-driven activation of p65/RelA enhances transcription of HuR [133], which in turn stabilizes *Eif4e* mRNA via binding to AU-rich elements within its 3′-UTR [134]. Conversely, cluster knockout-induced up-regulation of PTEN inhibits AKT pathway, thus destabilizing *eIF4E* mRNA, which, together with the reduced mTOR-dependent phosphorylation of its product, leads to impaired protein synthesis, including TH, DDC and DAT.

In addition to eIF4E, the miR-292a knockout cells revealed down-regulation of another translation initiation factor, the p97 isoform of eIF4G2. This may reflect a caspase-dependent cleavage of this protein towards an active IRES-dependent p86 isoform in response to apoptosis [135]. Our *in vitro* FUNCAT experiments directly demonstrate that miR-292a loss impairs translation initiation rather than elongation. Using the methionine analogue L-azidohomoalanine to label nascent proteins, we found that an elongation inhibitor anisomycin further suppressed protein synthesis in miR-292a-deficient MN9D cells, indicating that elongation was at least partially intact in the system. In contrast, lactimidomycin failed to exert additional inhibitory effects, indicating that the early translocation step targeted by this specific translation initiation inhibitor is already impaired in these cells. This pattern sharply contrasted with untargeted MN9D cells, where both inhibitors effectively suppressed translation. Altogether, these data indicate that miR-292a maintains the early steps of translation initiation. Accordingly, we detected impaired protein synthesis both in the striatum and SN^DA^ neurons of SN^DA^:miR-290-295CKO mice as early as 2 months after transduction, in agreement with our previous data reporting that mature DA neurons in mice with a deletion of PTEN in these cells reveal a prominent up-regulation of mTORC1, which activates the translation regulators eIF4B, eIF4E and S6 ribosomal protein [91]. In that same study, we have also reported that such boost of the PI3K-AKT-mTORC1-protein synthesis pathways in mature DA neurons protects them from degeneration and elevates the levels of proteins critical for DA biogenesis, such as TH and DAT [91].

Importantly, despite *Pten* was predicted to be the only common target of all cluster microRNAs within the thirty mostly up-regulated genes in SN^DA^:miR-290-295CKO animals, we may not include a critical contribution of other targets and, accordingly, other pathways apart from the PI3K-AKT-mTORC1-protein synthesis, in line with the typical multi-target mechanism of these small non-coding RNAs. Indeed, suppression of mRNA levels of *Eif4e*, *Th*, *Ddc* and *Dat* in isolated mature SN^DA^ neurons of knockout mice suggests to an even more complex cytoprotective role of these microRNAs regulating transcription and/or stability of these and other genes. Transcription of both *Th* and *Dat* is governed by availability of the critical dopaminergic transcription factor NURR1 [136]. Stability of the latter is tightly controlled by the phosphorylation state. Whereas α-synuclein-activated GSK-3β phosphorylates residues within the 123-134 residues of NURR1 to trigger its ubiquitination and proteasomal degradation [137], AKT phosphorylates GSK-3β at Ser9 to inhibit its activity and thereby stabilize NURR1 [138]. Conversely, NURR1 destabilization in response to PTEN-induced inhibition of the PI3K-AKT pathways in SN^DA^ neurons of miR-290-295 animals leads to early onset transcriptional silencing of both *Th* and *Dat* promoters, which, in addition to translation impairment, further contributes to a down-regulation of these proteins critical for DA biogenesis.

### Limitations of the study

The immortalized murine dopaminergic neuronal MN9D model used throughout this study is imperfect in many ways, primarily due to hybridization with N18TG2 neuroblastoma cells. However, as brilliantly stated by George E. P. Box, “All models are wrong, but some are useful”. While primary or induced pluripotent stem cells (iPSC)-derived neurons would not resolve the developmental vs. mature stage problem rendering them to be imperfect models, it’s also technically more difficult to propagate them, as well as modify their genome and epigenome. Instead, we chose to validate the mechanistic *in vitro* insights directly in mice.

Since our study was focused on the role of miR-290-295 in the adult midbrain DA neurons, we have not studied whether these stem cell-specific microRNAs remained abundant throughout developmental stages of this lineage or reemerged at later time points of the embryogenesis. Similarly, we haven’t characterized the roles of this cluster throughout development. As such, we have not quantified the numbers of DA neurons within the developing mesencephalon. This and other highly relevant objectives warrant further investigations.

### Conclusion

Altogether, our study revealed a previously unrecognized role of the *Pten*-targeting cluster miR-290-295, the most SN^DA^-enriched microRNAs within the SNpc of adult mice, and particularly miR-292a-3p, in maintaining mature DA biogenesis and protein synthesis, as well as protecting mice from locomotor deficits and other hallmarks of neurodegeneration. These data expand our understanding of post-transcriptional regulation in mature neurons by stem cell-specific non-coding RNAs, which may open a new page in age-related DA neuron vulnerability and neurodegenerative diseases, such as PD.

## METHODS

The detailed materials and methods, as well as the **Source data** for FACS gating strategy, TaqMan profiling and western blot experiments are available as Supplementary Data at NSR online.

## ACKNOWLEDGEMENTS

We thank Yun Tang for help with animal maintenance, Heike Alter for technical assistance, Qingxuan Lai, Günther Schütz and Yan Zhang for support, as well as Jinchang Li for assistance with image analysis.

## AUTHOR CONTRIBUTIONS

Conceptualization: IAV

Methodology: ZL, YX, NM, HJ, YL, XK, DS, AD, WZ, IAV

Investigation: ZL, YX, NM, RL, ET, NK, DSN, YL, XK, AD, IAV

Visualization: ZL, IAV

Funding acquisition: WZ, IAV

Project administration: WZ, IAV

Supervision: IAV

Writing – original draft: ZL, IAV

Writing – review & editing: ZL, AD, IAV

All authors reviewed and contributed to the manuscript.

### Data and materials availability

All data are available in the main text or the supplementary data.

## Supplementary Data

EXTENDED MATERIALS AND METHODS

SUPPLEMENTARY REFERENCES

Figures S1 to S4

Tables S1 to S3

Source data

Videos S1 to S5

