## Supplementary Data for "Selective abundance of miR-290-295 in the adult *Substantia Nigra* dopamine neurons is neuroprotective via preservation of protein synthesis"

**The PDF file includes:**

EXTENDED MATERIALS AND METHODS
SUPPLEMENTARY REFERENCES
Figures S1 to S4
Tables S1 to S3

**Other Supplementary data for this manuscript includes the following:**

Videos S1 to S5
Source data

### EXTENDED MATERIALS AND METHODS

#### *Animal models*

All experimental procedures were conducted in compliance with Animal Research: Reporting of *In Vivo* Experiments (ARRIVE) guidelines in the German Cancer Research Center (DKFZ) or Shanghai Jiao Tong University and approved by the institutional and local authorities of Germany (Z058I02, 35-9185.82/A-19/02, 35-9185.81/G-180/08, 35-9185.81/G-172/10 and 35-9185.81/G-154/10) or China (A2021092 and A2024195), respectively. Mice of both genders were maintained on the C57BL/6 genetic background (at least nine backcrosses) with a 12/12 h light/dark cycle, free access to water and standard (Kliba Nafag, #3437 and Jiangsu Xietong Biological Technology, #1010088) or methionine-deficient chow (Jiangsu Xietong Biological Technology, #XT93G). Please find details and ages of all groups in figure legends and **Table S1**, unless otherwise stated.

Dicer1$^{tm1Mmk}$ [1] (MGI:3835856) or B6.129S4-*Pten*$^{tm1Hwu}$/J [2] (Jackson lab, #006440) or *Rrn3*$^{tm1.1Igt}$/TmssJ [3] (Jackson lab, #039549) lines were crossed with Tg(Slc6a3-cre/ERT2)1Span mice [4] (MGI:3578093) to generate Dat$^{Cre\text{-}ERT2}$:Dicer1$^{fl/fl}$, Dat$^{Cre\text{-}ERT2}$:Pten$^{fl/fl}$ and Dat$^{Cre\text{-}ERT2}$:Rrn3$^{fl/fl}$ mice, respectively. Dat$^{Cre\text{-}ERT2}$:Rrn3$^{fl/fl}$Pten$^{fl/fl}$ mice were generated by intercrossing the latter two mouse lines. Temporal control to initiate recombination at specific time in the adulthood was achieved by nuclear translocation of Cre-ERT2 recombinase induced by intraperitoneal (i. p.) injections of 1 mg tamoxifen (Sigma-Aldrich, #T5648) diluted in sunflower oil twice a day for 5 days as described previously [5]. Dopamine transporter (DAT) promoter ensured expression of Cre in dopamine neurons.

B6;129S4-*Mirc5*$^{tm1Jae}$/J [6] (Jackson lab, #017337) line generated miR-290-295$^{+/-}$ and miR-290-295$^{-/-}$ mice. B6;129-*Gt(ROSA)26Sor*$^{tm1(CAG\text{-}cas9*,\text{-}EGFP)Fezh}$/J [7] and B6.SJL-*Slc6a3*$^{tm1.1(cre)Bkmn}$/J [8] (Jackson lab, #024857 and #006660, respectively) lines were crossed to generate Dat$^{Cre}$:SpCas9-GFP mice. *Slc6a3*$^{em(Cre)}$ mice (MGI:94862) on the C57BL/6J background were purchased from GENEPAX (#GAP1037) and used for Dat-driven overexpression and SaCas9-dependent knockout.

Since group allocation was based on the genotyping results, no randomization techniques were used. All cages were re-arranged randomly twice per week during routine bedding changes. During experimental procedures, animals from different cages were tested in the order from left to right and from top to bottom; for analyses, animals in the same cage were picked in a random order. Male mice were always tested before females. To prevent environmental differences, experimental or control mice were housed together in the same animal room from the birth to death and the behavior tests were performed in the same dedicated room for consistency. During testing, researchers randomly selected animals and assigned temporary visual markers, such as colored pen marks or tail line patterns, without knowledge of group identity. Group allocation was only revealed post-assessment by checking individual animal IDs, followed by data analysis. Definition of outcome measures for each experiment are outlined in the figure legends and the main text.

Mice were euthanized, striata and midbrain tissues were dissected from the 3$^{rd}$ to 6$^{th}$ and from the 7$^{th}$ to 10$^{th}$ rostral planes, respectively, by 1 mm coronal brain matrix (RWD Life Science, #68707) in 1 × phosphate-buffered saline (PBS) on ice. For RNA analyses, all reagents were treated with diethyl pyrocarbonate (DEPC). Fixed in 4% PFA tissues were cut to 6 µm-thick paraffin sections or 20 µm-thick cryoslices according to standard procedures. Striatal tissues were snap-frozen and stored at -80 °C for striatal DA content analysis by the DA high sensitive ELISA Kit (ImmuSmol, #BA E-5300). For western blot analysis, tissues were snap-frozen and stored at -80 °C prior to protein isolation.

#### *Fluorescent labeling and isolation of $SN^{DA}$ neurons*

As described previously [5], at least seven days before isolation, *Substantia Nigra pars compacta* (SNpc) dopamine ($SN^{DA}$) neurons were labeled fluorescently by retrobeads (Lumafluor) stereotaxically injected into striata of wild type mice and tamoxifen-induced $Dat^{Cre\text{-}ERT2}$:$Dicer1^{fl/fl}$ or control littermates (**Table S1**), followed by a direct transfer of the dissected brains to Ambion™ RNA*later*™ (Invitrogen, #AM7020) and storage according to the manufacturer's instructions. The details about laser-assisted microdissection of individual $SN^{DA}$ neurons and subsequent RNA isolation from wild type mice were described previously [5]. For dissection of SNpc by punching, we first sliced the RNA*later*™-submerged brains from $Dat^{Cre\text{-}ERT2}$:$Dicer1^{fl/fl}$ or control littermates by a vibratome, collecting 2 or 3 400 µm-thick sections per RNA*later*™-containing Eppendorf tube. SNpc patches were then dissected from sections mounted on SuperFrost slides under the cell^R fluorescent microscope (Olympus) using a 0.75 mm Harris Uni-Core punching tool (Plano GMBH, #15072) with a tip squeezed with pliers to an oval form to outline a typical for SNpc grouping of fluorescently labelled $SN^{DA}$ neurons, as depicted in **Figure 1A**. Patches collected in RNA*later*™-containing 2 ml Eppendorf tubes were then processed for total RNA isolation using the miRNeasy Mini kit buffers (Qiagen, #217004), but instead of the columns provided in the kit, the RNeasy MinElute columns (Qiagen, #74204) were used to reduce the elution volume. Super-Script III first-strand synthesis kit (Invitrogen, #18080051) was used for cDNA synthesis.

#### *Cell culture*

Mouse dopaminergic neuronal MN9D (BeiNa Culture Collection, #BNCC379310, a generous gift from Maozhong Sun), human embryonic kidney HEK293T and mouse hippocampal HT-22 (both from Cell Bank of Shanghai Institutes for Biological Sciences) immortalized cells were cultured at 37 °C with 5% of $CO_2$ in Dulbecco's Modified Eagle Medium (DMEM) with 10% fetal bovine serum (FBS) using standard cell culture techniques (Gibco, Thermo Fisher Scientific, #C11995500BT and #10099-141, respectively). Viability of intact and miR-292a-depleted MN9D cells with or without Lipofectamine 2000 (Thermo Fischer Scientific, #11668019)-assisted transfection with 30 nM double-stranded miR-292a-3p mimic (GenePharma, #B02001-134387), 800 nM siRNA targeting *Pten* (Ribobio Technology, #siG1177170201-1-5) or non-targeting controls (referred to as NC, GenePharma, #B02001-1111 and Ribobio Technology, #siN05815122147, respectively) for 48 hours, was assessed by the Cell counting kit 8 (CCK8, Sangon Biotech, #E606335-0500) at 450 nm according to the standard protocol. Total RNA from MN9D cells was isolated by RNA Easy Fast Tissue/Cell Kit (Tiangen Biotech, #4992732) or MolPure® Cell RNA Kit (Yeasen Biotechnology, #19231ES50) followed by cDNA synthesis by PrimeScript RT reagent Kit with gDNA Eraser (Takara Bio, #RR047A).

DIANA microT-CDS V 5.0 with the miTG score set to 0.5 was used to predict the targets of miR-290-295 [9]. The intact and mutated (synthesized by Tsingke Biotechnology) *Pten* 3′-UTR fragments were subcloned into pmirGLO plasmid (Promega, #E1330) by In-Fusion HD Cloning Kit (Takara Bio, #639648) and transfected using Lipofectamine 2000 (Thermo Fisher Scientific, #11668019) together with double-stranded miR-290a-3p, miR-291a-3p, miR-292a-3p, miR-293-3p, miR-294-3p or NC mimics (GenePharma, #B02001-65298, -64844, -134387, -65299, -64842 and -1111, respectively) in HEK293T cells (see **Table S2** and **Figures 5F and 5G**). Dual-luciferase reporter assay was conducted according to the manufacturer's instructions (Promega, #E1910) as described previously [10].

#### *Vectors*

Guide RNAs (gRNAs) with the highest on-target and lowest off-target activities were predicted by the CHOPCHOP tool (http://chopchop.cbu.uib.no/) [11] (**Table S2**). On-target activity of SpCas9 and SaCas9 gRNAs was validated *in vitro* by the Noodles system, as previously described [12], or in MN9D cells, respectively. Off-target thresholds for SpCas9 and SaCas9 gRNAs was set to 3 and 2, respectively, due to a lower overall off-target activity of the SaCas9 system [13]. Next, we used Lipofectamine 2000 (Thermo Fischer Scientific, #11668019) to transfect HT-22 cells with HP180 plasmid (a generous gift from Hui Yang) equipped with BbsI-HF (NEW ENGLAND BIOLABS, #R3539S) enzyme-subcloned gRNAs to extract total genomic DNA for subsequent PCR with primers spanning the off-target sites predicted by CHOPCHOP (**Table S2**). The amplified products of the expected lengths were then gel-extracted for evaluation of off-target activities of each gRNA by next-generation Illumina NovaSeq (Genewiz), as described previously [10] (**Table S3**).

For HP180.2 construction (**Source data**), the previously designed HP180.3 plasmid containing three gRNA cassettes [14], was opened by BaeI (NEW ENGLAND BIOLABS, #R0613S) and BbsI-HF using 1× CutSmart buffer (NEW ENGLAND BIOLABS, #B7204S). The resulting 10,640 bp backbone with the 5′ - GTTT and TTCCC - 3′ overhangs was then ligated with the annealed and phosphorylated oligonucleotides 5′ - CAC CGG GTC TTC GAG AAG ACC T - 3′ and 5′ - AAA CAG GTC TCC TCG AAG ACC CCG TGG GGA A - 3′ by Quick ligase (NEW ENGLAND BIOLABS, #M2200S).

To generate DA:miR-292aCKO and $SN^{DA}$:miR-290-295CKO models, two gRNAs were subcloned to each AAV-gRNA-mCherry vector using SapI (NEW ENGLAND BIOLABS, #R0569S) and BaeI as described previously [10]. The resulting constructs were packaged into PHP.eB or rAAV-9 serotypes (WZBiosciences) to conditionally knockout miR-292a or miR-290-295 in mature DA neurons by respectively tail vein or stereotaxic injections into SNpc of $Dat^{Cre}$:SpCas9-GFP mice (**Tables S1-S2**). To generate $SN^{DA}$:miR-290-295CKO$_{SaCas9}$ model, SaCas9-gRNA$_{290}$ and SaCas9-gRNA$_{295}$ were subcloned into an empty control elongation factor 1α (EFS)$_{promoter}$-driven construct to generate the EFS$_{promoter}$-DIO-SaCas9-U6-gRNA$_{290}$-U6-gRNA$_{295}$ vector (BrainVTA). For miR-292a overexpression, the hairpin structure of miR-292a was subcloned into the 5′-untranslated region (UTR) of tdTomato within the control human synapsin (hSyn)$_{promoter}$-driven construct to generate hSyn$_{promoter}$-DIO-miR-292a-tdTomato vector (BrainVTA). All four Cre-dependent rAAV vectors were packaged into neuron-specific CAP.B10 capsids and used for stereotaxic injections into the SNpc of adult *Slc6a3*$^{em(Cre)}$ mice to achieve a knock-out or overexpression of the cluster or miR-292a, respectively (**Tables S1-S2**). Such CRISPR-Cas9-based knockout approaches provided a temporal control, so that deletion occurred only upon injection of rAAV encoding the relevant gRNA, allowing targeting the genes in Cre-expressing neurons. Stereotaxically injected rAAV-9 or rAAV-CAP.B10 allowed high efficiency of neuronal transduction, together with the spatial control to target SN. Tail vein injection of blood-brain barrier-permeable rAAV-Php.eB allowed less invasive and less brain-damaging approach to target the neurons.

#### *Fluorescence activated cell sorting and flow cytometry analysis*

For construction of miR-292aKO clones, one gRNA per each strand of microRNA (**Table S2**) was subcloned to the GFP-equipped HP180.2 plasmid (**Source data** and see above) followed by its transfection to MN9D cells by Lipofectamine 2000 (Thermo Fischer Scientific, #11668019) and fluorescence-activated cell sorting (FACS) for $GFP^+$ cells by FACSAria II flow cytometer (BD). Single cells were collected in 96-well plate, propagated and sampled (Accurate Biology, #AG21009) for genomic DNA Sanger sequencing (Genewiz) to detect on-target mutations. Next, miR-292a-3p expression in selected clones was validated by real-time quantitative reverse transcription polymerase chain reaction (qRT-PCR).

For FACS-isolation of $SN^{DA}$ neurons, the brain samples were prepared as previously described [15]. Briefly, sagittally halved ventral midbrain from the rostral $7^{th}$ to $10^{th}$ planes were dissected by 1 mm coronal brain matrix (RWD Life Science, #68707) in cold 1 × PBS-DEPC on ice, followed by separation of the SNpc from the VTA according to the protocol published elsewhere [16] to only collect $SN^{DA}$ neurons. SNpc tissue was minced in cold 1 × PBS-DEPC on ice and digested with hibernate™-E medium containing 2 mg/mL collagenase IV (Gibco, Thermo Fisher Scientific, #A1247601 and #17104-019, respectively) at 37 °C for 15 min. After centrifugation at 150 g for 3 min at RT, the pellet was resuspended in hibernate™-E medium with 1 mg/ml papain (Shanghai Yuanye Bio-Technology, #S10011). 20 min after incubation at 37 °C, the digestion was stopped by addition of hibernate E medium and passing the sample through a 40-µm cell strainer. Cells were centrifuged at 500 g for 5 min at 4 °C, the pellet was resuspended in 100 µL 1 × PBS containing 1 µL FITC-conjugated DAT antibody (Alomone labs, #AMT-003-F) and incubated on ice in the dark for 25 min, followed by the addition of 2 mL 1 × PBS. After another centrifugation at 500 × g for 5 min at 4 °C, the pellet was resuspended in 1 × PBS containing 1% FBS (Gibco, Thermo Fisher Scientific, #10099-141). The cell suspension was kept on ice for subsequent FACS isolation using a FACSAria II flow cytometer (BD). Of note, the antibody incubation step was not performed if DA neurons carried endogenous GFP labeling. First, live and single cells were gated using Forward Scatter-Area (FSC-A) and Side Scatter-Area (SSC-A) to exclude cell fragments, aggregates, and dead cells. Gate 1 was set around the main cell population (typically located in the high-intensity FSC-A region), thereby excluding events with high FSC-A and high SSC-A, which are associated with noise or aggregates. Next, the presence of single versus multiple cells passing through the detector was evaluated by comparing FSC-Width (FSC-W) and FSC-A or FSC-Height (FSC-H) and FSC-A. For single cells, FSC-W/FSC-H was approximately equal to FSC-A, while for aggregates, FSC-W/FSC-H was greater than FSC-A. Gate 2 was applied to exclude events where FSC-A > FSC-W (i.e., non-single cells), retaining only the major peaks where FSC-A ≈ FSC-W. To further ensure the selection of single cells, a cross-validation step was performed by analyzing the Side Scatter Width (SSC-W) vs. Side Scatter Area (SSC-A). Events with SSC-W > SSC-A were considered doublets and excluded by Gate 3, thus retaining only true single cells. Finally, using lasers at 488 nm and/or 633 nm, fluorescently labeled cells were identified. A two-dimensional quadrant gate was applied, with the four quadrants (Q1-Q4) representing the following combinations: Q1, $GFP^{-}mCherry^{+}$; Q2: $GFP^{+}mCherry^{+}$; Q3, $GFP^{-}mCherry^{-}$; Q4, $GFP^{+}mCherry^{-}$. A subset gate was subsequently set within Q2 to sort the target cells expressing both GFP and mCherry. The sorting was performed using a 100 µm nozzle, in 1 × PBS buffer, in Purify mode (high purity mode). Fluorescence compensation was performed prior to sorting using $GFP^{-}mCherry^{+}$ and $GFP^{+}mCherry^{-}$ control samples to correct for spectral overlap. Sorted cells were collected into 200 µL V-shaped RNase-free microcentrifuge tubes containing lysis buffer. Subsequent downstream applications included microRNA and mRNA expression assays or Smart-seq2 library preparation.

For measurement of reactive oxygen species (ROS) levels in MN9D cells, H2DCFDA (MedChemExpress, #HY-D0940) was added to cell cultures at a final concentration of 5 µM. Following a 15 min incubation, cells were washed with 1 × PBS and harvested via trypsinization. Cells were resuspended in 1 × PBS prior to flow cytometric analysis.

For intracellular cytoplasmic protein staining, cells were collected by trypsin, fixed in 4% PFA for 15 min on ice, and permeabilized with 0.25% Triton X-100 for 15 min on ice. Cells were then incubated on ice for 25 min with primary antibodies, including rabbit anti-PTEN, rabbit anti-eIF4E and rabbit anti-eIF4G2/p97 (ABclonal, 1:330 #A11193, 1:330

#A22681 and 1:300 #A21193, respectively), followed by the incubation with CoraLite488-conjugated goat anti-rabbit IgG (Proteintech, 1:330 #SA00013-2) for 25 min on ice in the dark. After two washes with 1 × PBS, samples were subjected to flow cytometric analysis.

For flow cytometry gating, cells were first gated based on FSC-A and SSC-A, followed by FSC-H and FSC-W. A non-fluorescent control sample was used for voltage adjustment.

#### *Smart-seq2 profiling*

For Smart-seq2 profiling (Genewiz), $SN^{DA}$ neurons were FACS-isolated from $SN^{DA}$:miR-290-295CKO mice (one 13-month-old female, one 9-month-old female and one 15-month-old male) and Control mice (one 15-month-old male and two 13-month-old males). 50 sorted $GFP^{+}mCherry^{+}$ cells per mouse were directly collected in the lysis buffer (Tiagen), followed by total RNA isolation, purification using HiPure RNA Pure Micro Kits (Magen, #R2144-02) and pre-amplification by Discover-sc WTA Kit V2 (Vazyme, #N711-01). The amplified products were then quality-controlled using the Invitrogen Qubit 3.0 fluorometer for quantification and the Agilent 2100 Bioanalyzer for assessing RNA integrity and fragment size distribution. Library was constructed by TruePrep DNA Library Prep Kit V2 for Illumina (Vazyme, #TD501). GRCm38.p6 mouse reference genome was used for analysis. Samples with mapping ratio > 50% were used for transcriptome sequencing on the Illumina Novaseq 6000 system.

#### *qRT-PCR*

High-throughput microRNA expression profiling from punching tool-dissected SNpc was conducted using TaqMan Array Rodent MicroRNA A and B Cards Set v3.0 (Applied Biosystems, Thermo Fisher Scientific, #4444909) on a 7900HT Fast Real-Time PCR System (Applied Biosystems, Thermo Fisher Scientific, #4351405) following the manufacturer's protocol, as described previously [5]. Amplification plots were analyzed using SDS 2.2 software (Applied Biosystems, Thermo Fisher Scientific) with the Automatic Baseline setting and Manual $C_T$ threshold set to 0.2, as recommended by the manufacturer. The $C_T$ values were exported to qBase+ software (Biogazelle NV) for further analysis. For analysis of microRNAs in young and aged wild type mice or $Dat^{Cre\text{-}ERT2}$:$Dicer1^{fl/fl}$ mice, qRT-PCR was run for 40 cycles followed by global normalization with snoRNA135 and snoRNA202 as described previously [5], where undetectable nucleic acid signals were assigned a $C_T$ value of 42 or 40, respectively. After centering, the $C_T$ values were used for linear regression analysis. Relative expression was quantified via the $2^{-\Delta\Delta CT}$ method relative to *U6*.

qRT-PCR analyses were performed with a CFX96 Real-Time System or LightCycler® 480 Instrument II (Bio-Rad Laboratories, Hercules, #788BR05025, and Hoffmann-La Roche, #05015243001, respectively) using Hieff qRT-PCR SYBR Green Master Mix (No Rox) or TaqMan assay (Yeasen Biotechnology, #11201ES08, and Applied Biosystems, Thermo Fisher Scientific, #4440040, respectively) according to the manufacturer's instructions.

Mature or pre-microRNAs were analyzed via the poly(A)-tailing method. Hifair miRNA 1st Strand cDNA Synthesis Kit (Yeasen Biotechnology, #11148ES50) and Hieff miRNA Universal qRT-PCR SYBR Master Mix (Yeasen Biotechnology, #11171ES08) were used for cDNA synthesis and qRT-PCR. The kit provided the universal reverse primer. The unique target-specific forward primers for pre- and mature microRNA detection are listed in **Table S2**.

FACS-isolated $SN^{DA}$ neurons were used as a template to synthesize cDNA by CellAmp Whole Transcriptome Amplification Kit (Real Time) Ver.2 (Takara Bio, #3734). For detecting the expression levels of mature miRNAs within miR-290-295, the first strand cDNA was synthesized using a specific stem-loop primer following standard qRT-PCR procedures (see

**Table S2**). PCR was run for 65 cycles. Samples with undetectable nucleic acid signals were assigned a $C_T$ value of 66 for subsequent data analysis.

For all other assays, *U6*, glyceraldehyde-3-phosphate dehydrogenase (*Gapdh*), hypoxanthine phosphoribosyltransferase 1 (*Hprt1*) or beta-2-microglobin (*B2m*) were used as reference genes for normalized expression (see figure legends for details). The relative mRNA levels were calculated as $2^{-\Delta\Delta CT}$ (see **Table S2** for details).

#### *Behavioral assays*

Hindlimb clasping test, as an indicator of motor impairment or neurological dysfunction [17, 18], was performed based on previously reported protocols [19, 20]. Typically, healthy animals tend to extend their hindlimbs away from their abdomen when suspended, in contrast to an abnormal response by retracting or "clasping" hindlimbs towards their body in case of neurodegenerative or other conditions altering neural or motor functions. Briefly, the hindlimb position of a mouse gently lifted by its tail was observed for 30 seconds scaling from 0 to 3 to rate the phenotypes following phenotypes: 0 = both hindlimbs were consistently splayed outward (away from the abdomen); 0.5 = only one hindlimb was retracted towards the abdomen once, lasting for less than 15 seconds; 1 = only one hindlimb was retracted once, lasting for longer than 15 seconds; 1.5 = both hindlimbs were partially retracted, with each episode lasting for less than 15 seconds; 2 = both hindlimbs were partially retracted, with at least one of the episodes lasting for longer than 15 seconds; 2.5 = both hindlimbs were completely retracted, with each episode lasting for less than 15 seconds; 3 = both hindlimbs were completely retracted, with at least one of the episodes lasting for longer than 15 seconds .

The motor performance, balance, forelimb and hindlimb coordination were measured by constant-speed rotarod (Ugo Basile, #47650) assay at 25 rpm and 35 rpm as previously described [21], with the only modification: to not penalize mice for the forward rotations.

For the pole test, mice were placed with their head downwards on the top of a wooden gauze-wrapped 50 cm high standing rod with a diameter of 1 cm as described elsewhere [22]. To familiarize with the rod in advance every mouse was trained to climb. The average time to climb down the rod from five trials was used for subsequent analysis.

For the apomorphine-induced rotations, mice received i. p. injection of apomorphine (MedChemExpress, #HY-12723) solution at a dose of 0.25 mg per gram of body weight. 10 min after injection, each mouse was placed in a 50 cm × 50 cm × 50 cm open field arena for a 5 min test session, during which clockwise (ipsiversive) and anticlockwise (contraversive) rotations were manually counted. Two parameters were calculated to evaluate rotational bias: Δ net rotations = number of full 360° contraversive - number of full 360° ipsiversive rotations, and rotation baseline index = (number of full 360° contraversive - ipsiversive rotations) / number of full 360° total rotations.

#### *Histological analyses*

Immunohistochemical staining was performed as previously described [21], using the following primary antibodies: sheep anti-TH, rabbit anti-TH, rabbit anti-DDC and rat anti-DAT (Millipore, Merck, 1:1,000 #AB1542, 1:500 #MAB152, 1:1,000 #AB1569 and 1:500 #MAB369, respectively), followed by the respective biotinylated secondary antibodies: rabbit anti-sheep, goat anti-rabbit and goat anti-rat (1:500, Vectorlabs, #BA-6000, #BA-1,000 and #BA-9400, respectively).

For quantification of neuronal numbers, every $10^{th}$ cryoslice or every $6^{th}$ paraffin section was stained and analyzed. TH-positive, dopa decarboxylase (DDC)-positive and DAT-positive dopaminergic neurons were quantified in anatomically matched sections of the SNpc. TH-positive and DDC-positive neurons were identified based on a clearly defined soma with cytoplasmic chromogenic staining and a visible or lightly stained nuclear area; a profile was counted as one neuron only when a discernible nucleus and a substantial portion of the somatic contour were present, whereas somatic fragments lacking a nuclear profile were excluded. DAT-positive neurons were identified using morphology-based criteria adapted for the membrane-localized nature of DAT, requiring a discernible and relatively complete somatic contour, brown DAT immunoreactivity outlining the soma, and a central or eccentric unstained "white hole" corresponding to the nuclear shadow. Thin, elongated DAT-positive profiles or amorphous DAB-positive aggregates lacking a somatic contour were excluded to avoid confusion with axonal segments from neighboring neurons cut within the same plane or background DAB deposition. When necessary, brightness and contrast were uniformly adjusted across the entire image to enhance contour visibility without altering staining distribution. For each marker, the sum of all counted neurons across the selected sections was used for subsequent analysis.

For densitometry analysis of projections in the *Substantia Nigra pars reticulata* or striatum, one or five areas of the same size were chosen randomly per slice per mouse to quantify the average area or optical density by ImageJ (Fiji), respectively. All images used for densitometry were acquired under identical imaging parameters, including light source intensity, exposure time, camera gain and background correction, to ensure comparability across animals and experimental groups.

Tyramide signal amplification (TSA) kit (Hunan Aifang Biotechnology, #AFIHC034) was used to co-localize TH (rabbit anti-TH, 1:1,000, Millipore, Merck, #AB152) with mCherry (mouse anti-mCherry, 1:200, ABclonal Technology, #AE002).

*In situ* hybridization was performed as previously described [23]. Briefly, frozen midbrain sections were hybridized with digoxigenin-labelled miR-292a-3p probe (miRCURY LNA miRNA Detection Probe, QIAGEN, #339111 YD00616409-BCF) at 61 °C overnight in the RNase-free incubator followed by overnight incubation with sheep anti-DIG-AP (1:1,500, Hoffmann-La Roche Ltd, #11093274910) and staining with BM-Purple (Hoffmann-La Roche Ltd, #11442074001) according to a standard procedure. The hybridization signal was quantified by optical density analysis using ImageJ (Fiji), with all images acquired under the same illumination and exposure parameters as described above for densitometry.

#### *Western blotting*

Proteins isolated by RIPA buffer supplemented with phosphatase and protease inhibitors (ABclonal Technology, #RM02998 and #RM02916, respectively) were assayed in a standard western blot using the following antibodies: rabbit anti-DDC, rabbit anti-TH (both 1:1,000, Millipore, Merck #AB1569 and #AB152, respectively), sheep anti-TH (1:100,000, Millipore, Merck #AB1542, respectively), rabbit anti-DAT (1:500, ABclonal Technology, #A25875), rabbit anti β-ACTIN (1:2,000, ABclonal Technology, #AC026), mouse anti GAPDH (1:10,000, Millipore, Merck #MAB374) and HRP-conjugated goat anti-rabbit or anti-mouse (1:5,000, Cell Signaling Technology, #7074 and 1:2,000, ABclonal Technology, #AS003, respectively) or donkey anti-sheep (1:10,000, Santa Cruz Biotechnology, #sc-2473). Densitometry of bands from images acquired in ChemiDoc Touch System (Bio-Rad Laboratories) or LAS-3000 imaging system (Fuji) was analyzed by Fiji (ImageJ) or ImageGauge software (Fuji) and normalized to the reference proteins.

#### *Nascent protein synthesis analysis*

Nascent protein synthesis was assessed by fluorescent non-canonical amino acid tagging (FUNCAT) approach based on the click reaction between L-azidohomoalanine and alkyne-647 according to the protocol published elsewhere [24-26].

To assess nascent protein synthesis *in vitro*, MN9D and F4 cells were cultured until reaching 80% confluence. Cells were rinsed once with pre-warmed 1 × PBS and incubated in

1 mL methionine-free medium (Thermo Fisher Scientific, #21013024) for 30 min in a cell culture incubator. L-azidohomoalanine (Thermo Fisher Scientific, #C10102) was then added to the medium at a final concentration of 1 mM, and cells were incubated for an additional 2 h. For the protein-synthesis-inhibited control group, anisomycin (MedChemExpress, #HY-18982, final concentration 40 μM) and lactimidomycin (MedChemExpress, #HY-18979, final concentration 10 μM) were supplemented to the culture, followed by a further 30 min incubation. Cells were collected by trypsin, fixed in 4% PFA for 15 min, and permeabilized with 0.25% Triton X-100 for 15 min. Fluorescent labeling of L-azidohomoalanine was performed using the Click-iT Cell Reaction Buffer Kit (Thermo Fisher Scientific, #C10269) according to the manufacturer's instructions. For flow cytometry analysis, cells were initially gated by FSC-A and SSC-A, followed by FSC-H and FSC-W. A non-fluorescent control sample was used for instrument voltage calibration.

For the *in vivo* analysis of nascent protein synthesis, $SN^{DA}$:miR-290-295CKO or Control mice were fed the methionine-deficient chow for 7 d prior to the i. p. injection of 100 μg/g L-azidohomoalanine. After 3-4 h, the mice were euthanized followed by brain isolation and fixation in 4% PFA for 3 days and cyoslicing as described above. The slices were blocked for 18 h at 4 °C in a blocking buffer containing 0.05% Triton X-100, 10% normal goat serum, and 5% sucrose in PBS, gently washed six times with 1 × PBS for 5 min and incubated for 18 h at 4 °C in a click buffer containing 0.2 mM Tris[(1-benzyl-1H-1,2,3-triazol-4-yl)methyl]amine (Sigma-Aldrich, #678937), 0.5 mM Tris-(2-carboxyethyl) phosphine, hydrochloride (Thermo Fisher Scientific, #T2556), 2 μM fluorescent Alexa Fluor 647 alkyne (Thermo Fisher Scientific, #A10278) and 0.2 mM $CuSO_4$ (Sigma-Aldrich, #451657). Following ten washing cycles for 5 min each in 1 × PBS containing 0.5 mM EDTA and 1% Tween-20, the standard immunofluorescence staining was performed to colocalize the newly synthesized proteins with DA neurons labeled by TH. Images were acquired using an Olympus IXplore SpinSR10 spinning-disk confocal microscope equipped with a 60× oil-immersion objective (NA = 1.30) in Z-stack mode with a 2 μm step size. TH-positive dopaminergic neurons and L-azidohomoalanine signals were captured using the 488-nm and 640-nm channels, respectively. For quantification of nascent protein synthesis, Z-stack images were processed by maximum intensity projection in ImageJ (Fiji). Strictly excluding DAPI-positive nuclei, regions of interest of uniform size were manually defined within the somata of TH-positive neurons, where the mean fluorescence intensity of the L-azidohomoalanine signal was measured. To confirm the L-azidohomoalanine incorporation and the extent of protein synthesis inhibition in the cluster-deficient mice compared to controls with altered translatoin, 100 μg/g anisomycin was administered i.p. to the translation-altered Control mice 1 h before the L-azidohomoalanine injection.

##### *Statistical analyses*

Data that met key assumptions for regression analysis were analyzed by one- or two-way ANOVA followed by Holm-Šídák post-hoc test, two-tailed unpaired Student's t-test, one- or two-tailed unpaired Mann Whitney tests, Kruskal-Wallis test followed by Dunn's multiple comparisons test, or Log-Rank test as indicated in figure legends. Residual plots were used to evaluate model fit and detect potential patterns or outliers, indicating whether residuals were randomly distributed. Homoscedasticity plots assessed the constancy of residual variance across predicted values to verify the assumption of equal variance. Quantile-quantile (Q-Q) plots were employed to check for normality of residuals by comparing observed quantiles with those expected under a normal distribution. The individual mouse was considered as the experimental unit within the studies. The sample sizes (exact value of n are clearly indicated in the figure legends and **Source data**) were not determined using a formal a priori calculation, but were based on practical constraints and ethical considerations related to animal use. No animals were excluded in this study, but some data points were excluded from some analyses due to sample availability. While sufficient biological replicates were available for the initial marker assessment, subsequent analyses of additional markers involved reduced sample numbers due to limited remaining material. These exclusions were based solely on sample availability, and not due to experimental failure or selection bias. No animals or experimental units were excluded arbitrarily, and all usable data were incorporated into respective analyses. Effect sizes have not been reported throughout the study. Data in the figures and in the text are expressed as a mean ± standard error of mean (SEM). Analyses were performed using GraphPad Prism (V 9.0) or R (V 4.4.2). The details of the statistical tests and exact *p*-values are presented in **Source data**.

#### Figure S1

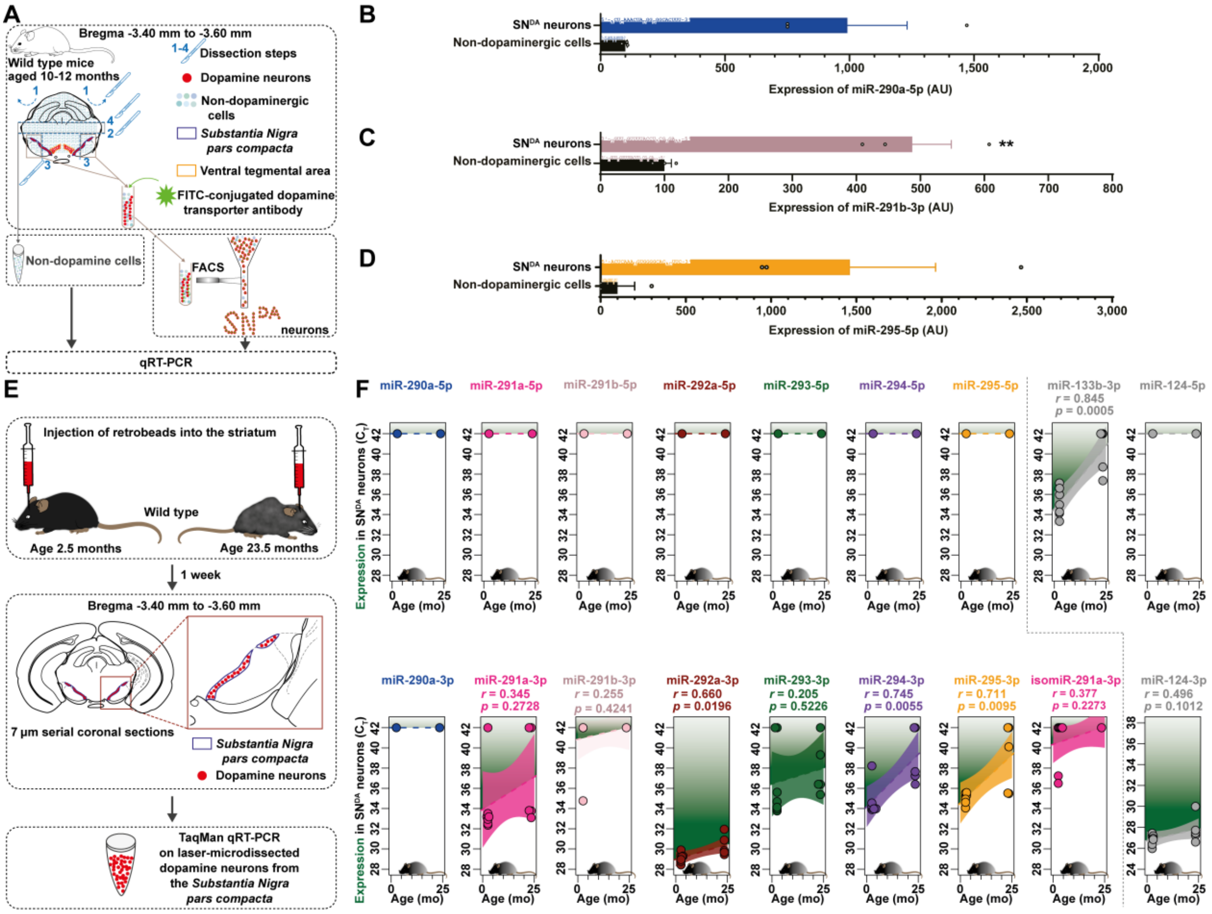


**Figure S1**. **Abundance of miR-290-295 in *Substantia Nigra* dopamine neurons.** (**A**) Experimental design of fluorescence-activated cell sorting (FACS) isolating *Substantia Nigra pars compacta* (SNpc) DA ($SN^{DA}$) neurons immunostained with dopamine transporter (DAT) antibody. For $SN^{DA}$ neurons, 6 *Slc6a3*$^{em(Cre)}$ mice were pooled into 3 biological replicates (2 mice each) and sorted by FACS. To obtain the non-dopaminergic cells, dorsal surrounding SNpc tissues were dissected from 3 *Slc6a3*$^{em(Cre)}$ male mice aged 10-12 months. (**B-D**) Expression of miR-290a-5p (**B**), miR-291b-3p (**C**) and miR-295-5p (**D**) in $SN^{DA}$ neurons versus non-dopaminergic cells (n = 3). (**E**) Experimental design of laser-assisted isolation of $SN^{DA}$ neurons from young (2.5 months old) and aged (23.5 months old) wild type male mice (n = 7 and 5, respectively). (**F**) Expression levels of indicated microRNAs and simple linear regression to evaluate age-dependent changes. Error bars represent SEM. **, $p < 0.01$ as analyzed by unpaired two-tailed Student's t-test. See the details about ages, sexes, FACS gating strategy and statistical methods in the **Source data**.

Figure S2

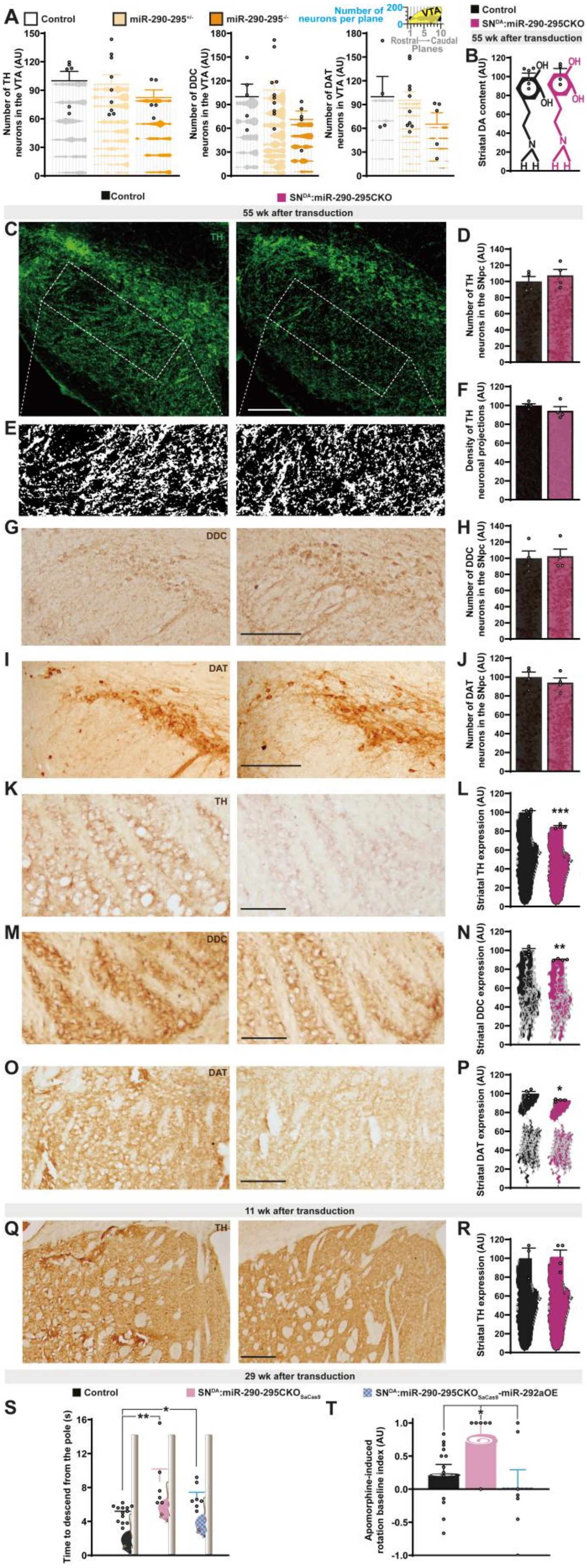


**Figure S2. Phenotypic and molecular characterization of mice deficient for miR-290-295.**

(**A**) Quantification of neurons immunostained for tyrosine hydroxylase (TH, **left**, n = 9, 5 and 6), dopa decarboxylase (DDC, **middle**, n = 10, 5 and 5) and DAT (**right**, n = 10, 5 and 4) in the ventral tegmental area (VTA) of miR-290-295$^{+/-}$, miR-290-295$^{-/-}$ and Control mice, respectively. (**B**) Striatal dopamine (DA) content in SN$^{DA}$:miR-290-295CKO and Control mice (n = 4 and 6, respectively). (**C-D**) Representative microphotographs (**C**) and quantification (**D**) of SN$^{DA}$ neurons immunostained for TH in SN$^{DA}$:miR-290-295CKO or Control mice (n = 4) 55 wk after transduction. (**E-F**) Visualization (**E**) and quantification of the area of TH-immunostained projections in the *Substantia Nigra pars reticulata* (**F**) of SN$^{DA}$:miR-290-295CKO or Control mice (n = 4) 55 wk after transduction. (**G-J**) Representative microphotographs (**G,I**) and quantification (**H,J**) of SN$^{DA}$ neurons immunostained for DDC (**G,H**) and DAT (**I,J**) in SN$^{DA}$:miR-290-295CKO or Control mice (n = 4) 55 wk after transduction. (**K-P**) Representative microphotographs (**K,M,O**) and quantification (**L,N,P**) of striatal tissues immunostained for TH (**K,L**), DDC (**M,N**) and DAT (**O,P**) in SN$^{DA}$:miR-290-295CKO or Control mice (n = 4) 55 wk after transduction. (**Q-R**) Representative microphotographs (**Q**) and quantification (**R**) of striatal TH content in SN$^{DA}$:miR-290-295CKO and Control mice 11 wk after transduction (n = 4 and 3, respectively). (**S**) Performance of SN$^{DA}$:miR-290-295CKO$_{SaCas9}$ and SN$^{DA}$:miR-290-295CKO$_{SaCas9}$-miR-292aOE and Control mice on the pole assays (n = 6, 7 and 13, respectively) 29 wk after transduction. (**T**) Apomorphine-induced rotation baseline index in SN$^{DA}$:miR-290-295CKO$_{SaCas9}$, SN$^{DA}$:miR-290-295CKO$_{SaCas9}$-miR-292aOE and Control mice (n = 6, 7 and 13, respectively) 29 wk after transduction. Rotation baseline index = (number of full 360° contraversive - ipsiversive rotations) / number of full 360° total rotations. Error bars represent SEM. ∗, $p < 0.05$; ∗∗, $p < 0.01$; ∗∗∗, $p < 0.001$ as analyzed by Kruskal-Wallis test with Dunn's multiple comparisons test (**A,S,T**), one-way ANOVA followed by post-hoc Holm-Šídák's test (**A**), or unpaired two-tailed Student's t-test (**L,N,P**). Scale bars in µm: 200 (**C,G,I,Q**), 50 (**K,M,O**). See the details about ages, sexes and statistical methods in the **Source data**.

#### Figure S3

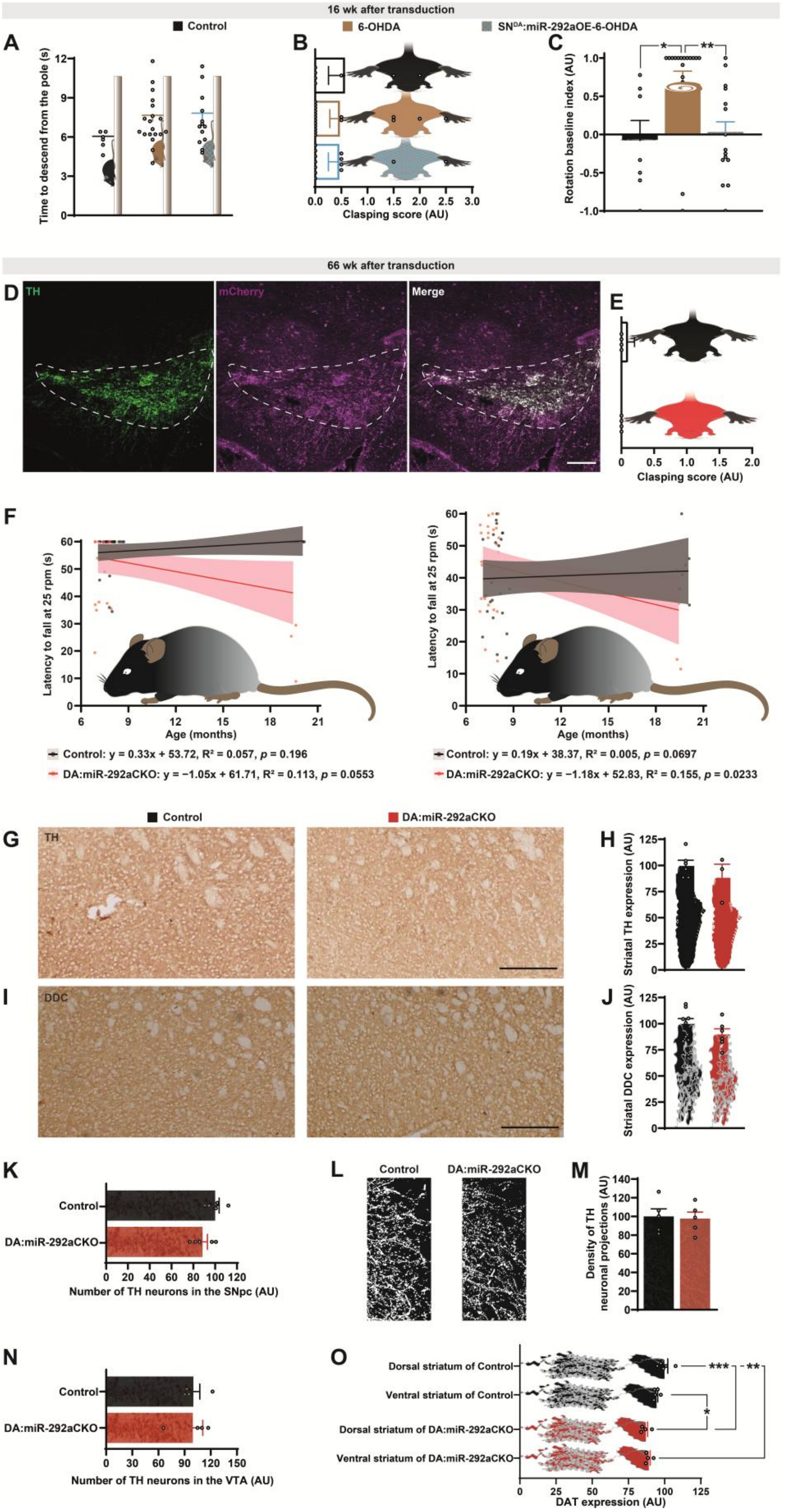


**Figure S3**. **Functional characterization of miR-292a in SN$^{DA}$ neurons.** (**A**) Time to descend from the pole in 6-OHDA, $SN^{DA}$:miR-292aOE-6-OHDA and Control mice (n = 19, 13 and 7, respectively) 16 wk after transduction. (**B**) Hindlimb clasping test score in 6-OHDA, $SN^{DA}$:miR-292aOE-6-OHDA and Control mice (n = 18, 13 and 7, respectively) 16 wk after transduction. (**C**) Apomorphine-induced rotation baseline index in 6-OHDA, $SN^{DA}$:miR-292aOE-6-OHDA and Control mice (n = 19, 15 and 7, respectively) 16 wk after transduction. Rotation baseline index = (number of full 360° contraversive - ipsiversive rotations) / number of full 360° total rotations. (**D**) Representative microphotographs of TH and mCherry localization in the ventral midbrain of DA:miR-292aCKO mice. (**E**) Hindlimb clasping test score in DA:miR-292aCKO and Control mice (n = 4 and 5, respectively) 66 wk after transduction. (**F**) Rotarod latency to fall was measured at 25 rpm (left) and 35 rpm (right) to evaluate age-associated motor performance by simple linear regression in DA:miR-292aCKO and Control mice. (**G-J**) Representative microphotographs (**G,I**) and quantification (**H,J**) of the striata immunostained for TH (**G,H,** n = 3 and 6, respectively) or DDC (**I,J,** n = 6 and 8, respectively) from DA:miR-292aCKO or Control mice 66 wk after transduction. (**K**) Numbers of TH-positive neurons in the SNpc of DA:miR-292aCKO or Control mice (n = 5) 66 wk after transduction. (**L-M**) Representative visualization (**L**) and quantification (**M**) of TH-immunostained projections in the *Substantia Nigra pars reticulata* of DA:miR-292aCKO or Control mice (n = 5) 66 wk after transduction. (**N**) Numbers of TH-positive neurons in the VTA of DA:miR-292aCKO or Control mice (n = 4) 66 wk after transduction. (**O**) DAT expression in the dorsal and ventral striatum of DA:miR-292aCKO or Control mice (n = 4 and 5, respectively) 66 wk after transduction. Error bars represent SEM. ∗, $p < 0.05$; ∗∗, $p < 0.01$; ∗∗∗, $p < 0.001$ as analyzed by Kruskal-Wallis test with Dunn's multiple comparisons test (**C**) or one-way ANOVA followed by post-hoc Holm-Šídák's test (**O**). Scale bars: 200 µm. See the details about ages, sexes and statistical methods in the **Source data**.

**Figure S4**

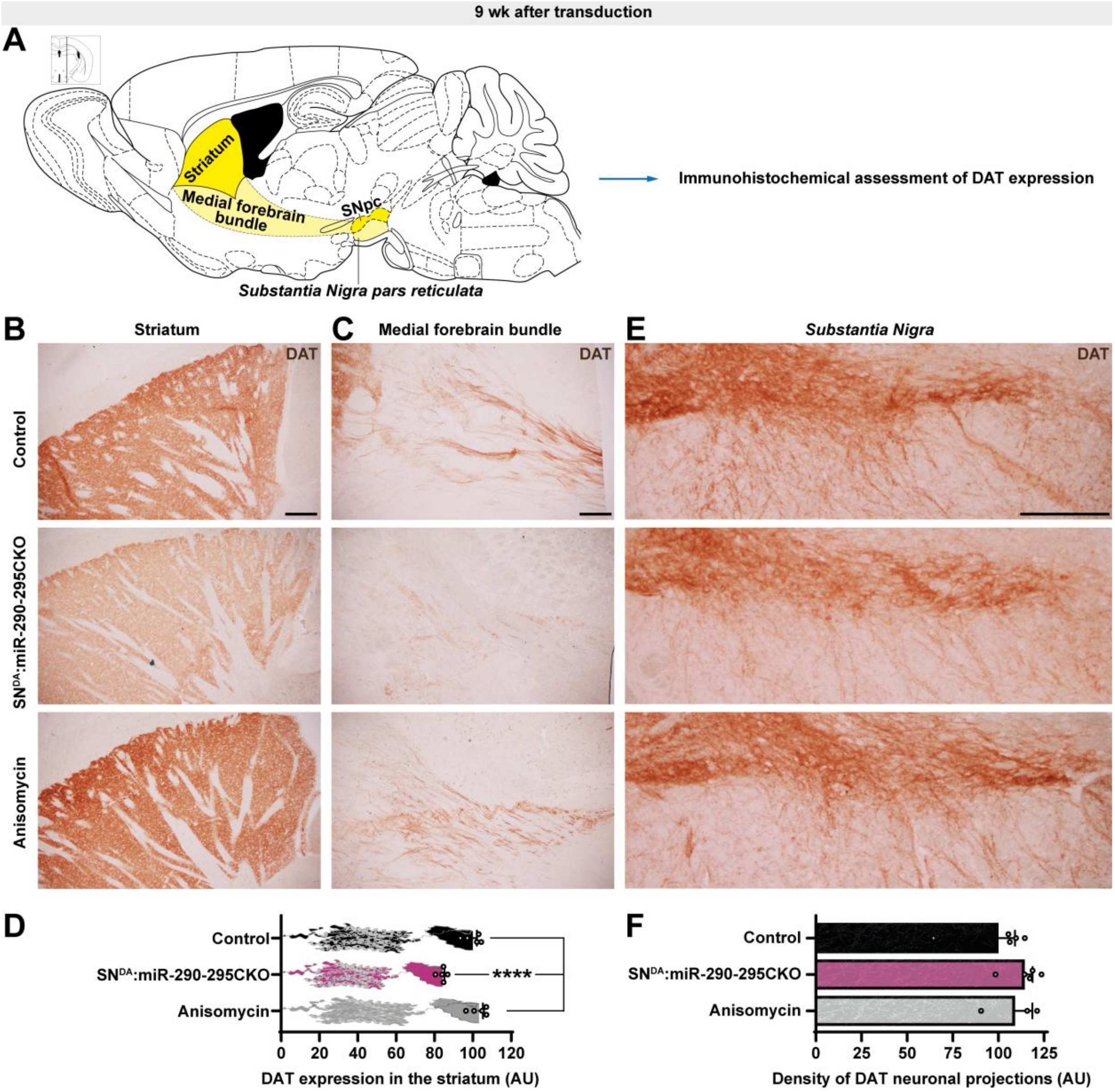


**Figure S4. Expression of DAT in mice with altered translation or SN$^{DA}$-specific knockout of miR-290-295.** (**A**) Anatomic location of the SNpc, the medial forebrain bundle and the striatum. (**B-F**) Representative microphotographs of DAT immunoreactivity in the striatum (**B**), medial forebrain bundle (**C**) or SNpc (**E**) and quantification of the striatal DAT expression (**D**, n = 5, 5 and 5, respectively) and the area of of DAT-immunostained projections in the *Substantia Nigra pars reticulata* (**F**, n = 5, 5 and 3, respectively) in SN$^{DA}$:miR-290-295CKO, Control and Anisomycin negative control mice 9 wk after transduction. Error bars represent SEM. ∗∗∗∗, $p < 0.0001$ as assessed by one-way ANOVA with post-hoc Holm-Šídák's test (**D**). Scale bar: 200 µm. See the details about ages, sexes and statistical methods in the **Source data**.

**Table S1. Details about stereotaxic and rAAV injections in this study.**

| Injected substances | Group | Details of the experimental groups† | Details of the control groups† | Injection site# and figure references |
|---|---|---|---|---|
| Red fluorescent retrobeads stereotaxically injected into the striatum | 1.1 | $Dat^{Cre\text{-}ERT2}$:$Dicer1^{fl/fl}$ mice injected intraperitoneally (i.p.) with 1 mg tamoxifen diluted in sunflower oil twice a day for 5 days followed by stereotaxic injection with 300 nL red fluorescent retrobeads per site. | $Dat^{Cre\text{-}ERT2}$:$Dicer1^{fl/fl}$ mice injected i.p. with sunflower oil twice a day for 5 days then stereotaxic injected with 300 nL red fluorescent retrobeads per site. | A/P +0.4 mm, M/L +/- 1.8 mm, D/V -3.5 mm (**Figure 1A-C**). |
| | 1.2 | Wild type mice injected with 300 nL red fluorescent retrobeads per site. | | A/P +0.4 mm, M/L +/- 1.8 mm, D/V -3.5 mm (**Figure 1E-F**). |
| | 1.3 | Wild type mice injected with 300 nL red fluorescent retrobeads per site. | | A/P +0.4 mm, M/L +/- 1.8 mm, D/V -3.5 mm (**Figure S1E-F**). |
| rAAV-9 equipped with $gRNA_{290/295}$ or empty constructs stereotaxically injected into the *Substantia Nigra pars compacta* | 2.1 | Homozygous $DAT^{Cre}$:SpCas9-GFP mice injected with 1 μL per site of rAAV ($7.72*10^{13}$ vg/mL). | Wild type mice injected with 1 μL per site of rAAV ($6.52*10^{13}$ vg/mL). | A/P -3, M/L ±1.2, D/V -4 and A/P -3.52, M/L ±1.3, D/V -3.9 (**Figures 2J-N** and **S2B-P**). |
| | 2.2 | $DAT^{Cre}$:SpCas9-GFP mice homozygous or heterozygous for both $DAT^{Cre}$ and SpCas9 injected with 1 μL per site of rAAV ($7.72*10^{13}$ vg/mL). | $DAT^{Cre}$:SpCas9-GFP mice homozygous or heterozygous for both $DAT^{Cre}$ and SpCas9 injected with 1 μL per site of rAAV ($6.52*10^{13}$ vg/mL). | A/P -3.08, M/L ±1.25, D/V -4.3 and A/P -3.52, M/L ±1.3, D/V -3.9 (**Figures 2J-L, O-P, 5A, D** and **S2Q-R**). |
| | 2.3 | $DAT^{Cre}$:SpCas9-GFP mice homozygous or heterozygous for $DAT^{Cre}$ and homozygous for SpCas9 injected with 1 μL per site of rAAV ($7.72*10^{13}$ vg/mL). | $DAT^{Cre}$:SpCas9-GFP mice homozygous or heterozygous for $DAT^{Cre}$ and homozygous for SpCas9 injected with 1 μL per site of rAAV ($6.52*10^{13}$ vg/mL). | A/P -3.08, M/L ±1.25, D/V -4.3 and A/P -3.52, M/L ±1.3, D/V -3.9 (**Figure 6H-L** and **S4**). |

<table>
<tr>
<td>rAAV-CAP.B10 equipped with SaCas9 and SaCas9-gRNA$_{290/295}$ and/or miR-292a or corresponding empty constructs stereotaxically injected to the Substantia Nigra pars compacta</td>
<td>3</td>
<td>Homozygous DAT$^{Cre}$ mice injected with 1 μL per site of rAAV mixtures (vol:vol = 1:1) following the below table:
<table>
<tr><th>Experimental group</th><th>Left hemisphere</th><th>Right hemisphere</th></tr>
<tr><td>SN$^{DA}$:miR-290-295CKO$_{SaCas9}$</td><td>rAAV-Empty (5.33*10$^{12}$ vg/mL) and rAAV-DIO-tdTomato (5.65*10$^{12}$ vg/mL)</td><td>rAAV-DIO-SaCas9-U6-gRNA$_{290}$-U6-gRNA$_{295}$ (5.09*10$^{12}$ vg/mL) and rAAV-DIO-tdTomato (5.65*10$^{12}$ vg/mL).</td></tr>
<tr><td>SN$^{DA}$:miR-290-295CKO$_{SaCas9}$-miR-292aOE</td><td>rAAV-Empty (5.33*10$^{12}$ vg/mL) and rAAV-DIO-tdTomato (5.65*10$^{12}$ vg/mL)</td><td>rAAV-DIO-SaCas9-U6-gRNA$_{290}$-U6-gRNA$_{295}$ (5.09*10$^{12}$ vg/mL) and rAAV-DIO-miR-292a-tdTomato (5.40*10$^{12}$ vg/mL).</td></tr>
</table>
</td>
<td>Homozygous DAT$^{Cre}$ mice injected with 1 μL per site of rAAV mixtures (vol:vol = 1:1) of rAAV-Empty (5.33*10$^{12}$ vg/mL) and rAAV-DIO-tdTomato (5.65*10$^{12}$ vg/mL) on both sides.</td>
<td>A/P -3.08, M/L ±1.25, D/V -4.3 and A/P -3.52, M/L ±1.3, D/V -3.9 (Figures 2Q-Z and S2S-T).</td>
</tr>
</table>

<table>
<tr>
<td>rAAV-CAP.B10 equipped with miR-292a or empty stereotaxically injected to the Substantia Nigra pars compacta and 6-OHDA or vehicle stereotaxically injected to the striatum</td>
<td>4</td>
<td>Homozygous $DAT^{Cre}$ mice or wilde type mice injected with 0.5 µL per site of rAAV diluted twice in saline in SNpc firstly following another injection of 6-OHDA or saline in the striatum following the below table:
<table>
<tr><th>Experimental group</th><th>Injection time</th><th>Left hemisphere</th><th>Right hemisphere</th></tr>
<tr><td rowspan="2">6-OHDA</td><td>$1^{st}$ injection</td><td>$DAT^{Cre}$ mice injected with rAAV-DIO-tdTomato ($5.65*10^{12}$ vg/mL) or wild type injected with rAAV-DIO-miR-292a-tdTomato ($5.40*10^{12}$ vg/mL).</td><td>$DAT^{Cre}$ mice injected with rAAV-DIO-tdTomato ($5.65*10^{12}$ vg/mL) or wild type injected with rAAV-DIO-miR-292a-tdTomato ($5.40*10^{12}$ vg/mL).</td></tr>
<tr><td>$2^{nd}$ injection</td><td>0.5 µL saline</td><td>0.5 µL 6-OHDA (3µg/µL)</td></tr>
<tr><td rowspan="2">$SN^{DA}$:miR-292aOE-6-OHDA</td><td>$1^{st}$ injection</td><td>$DAT^{Cre}$ mice injected with rAAV-DIO-miR-292a-tdTomato ($5.40*10^{12}$ vg/mL).</td><td>$DAT^{Cre}$ mice injected with rAAV-DIO-miR-292a-tdTomato ($5.40*10^{12}$ vg/mL).</td></tr>
<tr><td>$2^{nd}$ injection</td><td>0.5 µL saline</td><td>0.5 µL 6-OHDA (3µg/µL)</td></tr>
</table>
</td>
<td>Wild type mice injected with 0.5 µL per site of rAAV-DIO-miR-292a-tdTomato ($5.40*10^{12}$ vg/mL) diluted twice in saline in SNpc firstly following another injection of 0.5 µL saline in the striatum on both sides.</td>
<td>
<table>
<tr><th>Injection time</th><th>Sites</th></tr>
<tr><td>$1^{st}$ injection</td><td>A/P -3.08, M/L ±1.25, D/V -4.3 and A/P -3.52, M/L ±1.3, D/V -3.9</td></tr>
<tr><td>$2^{nd}$ injection</td><td>A/P +0.4 mm, M/L +/-1.8 mm, D/V -3.5 mm</td></tr>
</table>
(Figures 4A-G and S3A-C)
</td>
</tr>
</table>

| | | | | |
|---|---|---|---|---|
| rAAV-PHP.eB equipped with $gRNA_{292a\text{-}5/3p}$ or empty injected into the tail vein | 5 | Homozygous $DAT^{Cre}$:Cas9-GFP mice injected with 100 µL rAAV ($2.60*10^{13}$ vg/mL). | Wild type mice injected with 1.2 µL rAAV ($1.15*10^{13}$ vg/mL) diluted 6 times in 1 × phosphate-buffered saline (PBS). | Tail vein injection (**Figures 3B-C, 4H-N** and **S3D-O**). |

†Including genotypes, injection volumes per site, titers of rAAVs (in vg/mL).
#Relative to Bregma (mm): A/P, anterio-posterior; M/L, medio-lateral; D/V, dorso-ventral.

**Table S2. Oligonucleotides used in this work.**

| Oligonucleotides, order # | Antisense strand or forward primer (5′-3′) | Sense strand or reverse primer (5′-3′) | Company |
|---|---|---|---|
| | | **TaqMan qRT-PCR probe (Figures 1B-C and S1F)** | |
| miR-124a-5p (#002197) | | CGUGUUCACAGCGGACCUUGAU† | Applied Biosystems |
| miR-124a-3p (#001182) | | UAAGGCACGCGGUGAAUGCC | Applied Biosystems |
| miR-133b-3p (#002247) | | UUUGGUCCCCUUCAACCAGCUA | Applied Biosystems |
| miR-139-3p (#002546) | | UGGAGACGCGGCCCUGUUGGAG | Applied Biosystems |
| miR-290a-5p (#002590) | | ACUCAAACUAUGGGGGCACUUU | Applied Biosystems |
| miR-290a-3p (#002591) | | AAAGUGCCGCCUAGUUUUAAGCCC | Applied Biosystems |
| miR-291a-5p (#001202) | | CAUCAAAGUGGAGGCCCUCUCU | Applied Biosystems |
| isomiR-291a-3p (#001135) | | AAAGUGCUUCCACUUUGUGUGCC | Applied Biosystems |
| miR-291a-3p (#002592) | | AAAGUGCUUCCACUUUGUGUGC | Applied Biosystems |
| miR-291b-5p (#002537) | | GAUCAAAGUGGAGGCCCUCUCC | Applied Biosystems |
| miR-291b-3p (#002538) | | AAAGUGCAUCCAUUUUGUUUGU | Applied Biosystems |
| miR-292a-5p (#001055) | | ACUCAAACUGGGGGCUCUUUUG | Applied Biosystems |
| miR-292a-3p (#002593) | | AAAGUGCCGCCAGGUUUUGAGUGU | Applied Biosystems |
| miR-293-5p (#002594) | | ACUCAAACUGUGUGACAUUUUG | Applied Biosystems |
| miR-293-3p (#001794) | | AGUGCCGCAGAGUUUGUAGUGU | Applied Biosystems |
| miR-294-5p (#002595) | | ACUCAAAAUGGAGGCCCUAUCU | Applied Biosystems |
| miR-294-3p (#001056) | | AAAGUGCUUCCCUUUUGUGUGU | Applied Biosystems |
| miR-295-5p (#002596) | | ACUCAAAUGUGGGGCACACUUC | Applied Biosystems |
| miR-295-3p (#000189) | | AAAGUGCUACUACUUUUGAGUCU | Applied Biosystems |
| miR-302a-3p (#000529) | | UAAGUGCUUCCAUGUUUUGGUGA | Applied Biosystems |
| miR-409-5p (#002331) | | AGGUUACCCGAGCAACUUUGCAU | Applied Biosystems |
| snoRNA135 (#001230) | | CTAAAATAGCTGGAATTACCGGCAGATTGGTAGTGGTGAG CCTATGGTTTTCTGAAG | Applied Biosystems |

| | | | |
|---|---|---|---|
| snoRNA202 (#001232) | GCTGTACTGACTTGATGAAAGTACTTTTGAACCCTTTTCCATCTGATG | | Applied Biosystems |
| U6 (#001973) | GTGCTCGCTTCGGCAGCACATATACTAAAATTGGAACGATACAGAGAAGATTAGCATGGCCCCTGCGCAAGGATGACACGCAAATTCGTGAAGCGTTCCATATTTT | | Applied Biosystems |
| | ***In situ* probe (Figure 4I-J)** | | |
| miR-292a-3p | *ACTCAAAACCTGGCGGCACTTT*-Digoxigenin† | | Qiagen |
| **Stem loop primer for synthesis the first strand cDNA of miRNA (Figures 1F, 2M,S, 3E, 4B and S1B-D)** | | | |
| SL-miR-290a-5p | GTCGTATCCAGTGCAGGGTCCGAGGTATTCGCACTGGATACGACAAAGTG | | Tsingke |
| SL-miR-290a-3p | GTCGTATCCAGTGCAGGGTCCGAGGTATTCGCACTGGATACGACGGGCTT | | Tsingke |
| SL-miR-291a-5p | GTCGTATCCAGTGCAGGGTCCGAGGTATTCGCACTGGATACGACAGAGAG | | Tsingke |
| SL-isomiR-291a-3p | GTCGTATCCAGTGCAGGGTCCGAGGTATTCGCACTGGATACGACGGCACA | | Tsingke |
| SL-miR-291a-3p | GTCGTATCCAGTGCAGGGTCCGAGGTATTCGCACTGGATACGACGCACAC | | Tsingke |
| SL-miR-291b-5p | GTCGTATCCAGTGCAGGGTCCGAGGTATTCGCACTGGATACGACGGAGAG | | Tsingke |
| SL-miR-291b-3p | GTCGTATCCAGTGCAGGGTCCGAGGTATTCGCACTGGATACGACACAAAC | | Tsingke |
| SL-miR-292a-5p | GTCGTATCCAGTGCAGGGTCCGAGGTATTCGCACTGGATACGACCAAAAG | | Tsingke |
| SL-miR-292a-3p | GTCGTATCCAGTGCAGGGTCCGAGGTATTCGCACTGGATACGACACACTC | | Tsingke |
| SL-miR-293-5p | GTCGTATCCAGTGCAGGGTCCGAGGTATTCGCACTGGATACGACCAAAAT | | Tsingke |
| SL-miR-293-3p | GTCGTATCCAGTGCAGGGTCCGAGGTATTCGCACTGGATACGACACACTA | | Tsingke |
| SL-miR-294-5p | GTCGTATCCAGTGCAGGGTCCGAGGTATTCGCACTGGATACGACAGATAG | | Tsingke |
| SL-miR-294-3p | GTCGTATCCAGTGCAGGGTCCGAGGTATTCGCACTGGATACGACACACAC | | Tsingke |
| SL-miR-295-5p | GTCGTATCCAGTGCAGGGTCCGAGGTATTCGCACTGGATACGACGAAGTG | | Tsingke |
| SL-miR-295-3p | GTCGTATCCAGTGCAGGGTCCGAGGTATTCGCACTGGATACGACAGACTC | | Tsingke |
| | **Transfection of microRNA mimics *in vitro* (Figures 3H, O-P, R-S and 5G-H)** | | |
| miR-290a-3p | gcuuaaaacuaggcggcacuuuuu† | aaagugccgccuaguuuuaagccc | GenePharma |
| miR-291a-3p | acacaaaguggaagcacuuuuu | aaagugcuuccacuuugugugc | GenePharma |
| miR-292a-3p | acucaaaaccuggcggcacuuuuu | aaagugccgccagguuuugagugu | GenePharma |
| miR-293-3p | acuacaaacucugcggcacuuu | agugccgcagaguuuguagugu | GenePharma |

| | | | |
|---|---|---|---|
| miR-294-3p | acacaaaagggaagcacuuuuu | aaagugcuucccuuuugugugu | GenePharma |
| Negative control | acgugacacguucggagaatt | uucuccgaacgugucacgutt | GenePharma |
| | **Primers for qRT-PCR (Figures 1F, 2M, R-S,V-X, 3E, M-P, 4B, 5I, K-L, N-O and S1B-D)** | | |
| miR-290a-5p | GTGCAGGGTCCGAGGTATT | CGCGACTCAAACTATGGGGG | Tsingke |
| miR-290a-3p | GTGCAGGGTCCGAGGTATT | CGAAAGTGCCGCCTAGT | Tsingke |
| miR-291a-5p | GTGCAGGGTCCGAGGTATT | GCATCAAAGTGGAGGCCCT | Tsingke |
| isomiR-291a-3p | GTGCAGGGTCCGAGGTATT | CGCGAAAGTGCTTCCACTT | Tsingke |
| miR-291a-3p | GTGCAGGGTCCGAGGTATT | CGCGAAAGTGCTTCCACTT | Tsingke |
| miR-291b-5p | GTGCAGGGTCCGAGGTATT | CGATCAAAGTGGAGGCCCT | Tsingke |
| miR-291b-3p | GTGCAGGGTCCGAGGTATT | CGCGAAAGTGCATCCATT | Tsingke |
| miR-292a-5p | GTGCAGGGTCCGAGGTATT | GACTCAAACTGGGGGCTCTT | Tsingke |
| miR-292a-3p | GTGCAGGGTCCGAGGTATT | CAAAGTGCCGCCAGGTT | Tsingke |
| miR-293-5p | GTGCAGGGTCCGAGGTATT | CGCGACTCAAACTGTGTGACA | Tsingke |
| miR-293-3p | GTGCAGGGTCCGAGGTATT | CGAGTGCCGCAGAGTTTGT | Tsingke |
| miR-294-5p | GTGCAGGGTCCGAGGTATT | CGACTCAAAATGGAGGCCCT | Tsingke |
| miR-294-3p | GTGCAGGGTCCGAGGTATT | CGCGAAAGTGCTTCCCTT | Tsingke |
| miR-295-5p | GTGCAGGGTCCGAGGTATT | GACTCAAATGTGGGGCACAC | Tsingke |
| miR-295-3p | GTGCAGGGTCCGAGGTATT | GCGCGCAAAGTGCTACTACT | Tsingke |
| P(A)-Tailing-miR-290a-5p | ACTCAAACTATGGGGGCACTTT | Universal Reverse Primer (YEASEN Biotechnology, #11148-D) | Tsingke |
| P(A)-Tailing-miR-291a-5p | CATCAAAGTGGAGGCCCTCTCT | Universal Reverse Primer (YEASEN Biotechnology, #11148-D) | Tsingke |
| P(A)-Tailing-miR-292a-5p | ACTCAAACTGGGGGCTCTTTTG | Universal Reverse Primer (YEASEN Biotechnology, #11148-D) | Tsingke |
| P(A)-Tailing-miR-292a-3p | AGTGCCGCCAGGTTTTGAG | Universal Reverse Primer (YEASEN Biotechnology, #11148-D) | Tsingke |
| P(A)-Tailing-miR-295-5p | GGGGAAAGTGCTACTACTTTTGAGTCT | Universal Reverse Primer (YEASEN Biotechnology, #11148-D) | Tsingke |
| *U6* | CTCGCTTCGGCAGCACA | AACGCTTCACGAATTTGCGT | Tsingke |

| | | | |
|---|---|---|---|
| *Pten* | TGGATTCGACTTAGACTTGACCT | GCGGTGTCATAATGTCTCTCAG | Tsingke |
| *Th* | CCAAGGTTCATTGGACGGC | CTCTCCTCGAATACCACAGCC | Tsingke |
| *Ddc* | AGCTGACTATCTGGATGGCAT | ACCCCTGGCATGATTATCTTCT | Tsingke |
| *Dat* | GGTGCTGATTGCCTTCTCCAGT | GACAACGAAGCCAGAGGAGAAG | Tsingke |
| *Hprt1* | TCAGTCAACGGGGGACATAAA | GGGGCTGTACTGCTTAACCAG | Tsingke |
| *B2m* | TTCTGGTGCTTGTCTCACTGA | CAGTATGTTCGGCTTCCCATTC | Tsingke |
| *Gapdh* | CATCACTGCCACCCAGAAGACTG | ATGCCAGTGAGCTTCCCGTTCAG | Tsingke |
| **Sp/SaCas9-gRNAs (Figures 2J-Z, 3, 4, 5A,D, S6H-L, S2B-T, S3 and S4)** | | | |
| gRNA$_{290}$ | ATCTTGCGGTACTCAAACTATGG | | Genewiz |
| gRNA$_{295}$ | ATCTTGGTGAGACTCAAATGTGG | | Genewiz |
| SaCas9-gRNA$_{290}$ | AAAAAAAGTGCCCCCATAGTTTGAGT | | BrainVTA |
| SaCas9-gRNA$_{295}$ | CATAGAAAGTGCTACTACTTTTGAGT | | BrainVTA |
| gRNA$_{292a\text{-}5p}$ | CCAGCCTGTGATACTCAAACTGG | | Genewiz |
| gRNA$_{292a\text{-}3p}$ | ATCGGAAGAAAAGTGCCGCCAGG | | Genewiz |
| **Hairpin structure subcloned in rAAV vector to overexpress the miR-292a (Figures 2Q, S-Z, 4A-G, S2S-T and S3A-C)** | | | |
| GTAAGTATCAAGGTTACAAGACAGGTAGGGCTCTGCGTTTGCTCCAGGTAGTCCGCTGCTCCCTTGGGCCTGGGCCCA<br>CTGACAGCCCTGGTGCCTCTGGCCGGCTGCACACCTCCTGGCGGGCAGCTGTGAAAGTGCCGCCAGGTTTTGAGTGTT<br>GTTCTGGCAATACCTGACACTCAAAACCTGGCGGCACTTTCACGGAGGCCTGCCCTGACTGCCCACGGTGCCGTGGCC<br>AAAGAGGATCTAAGGGCACCGCTGAGGGCCTACCTAACCATCGTGGGGAATAAGGACAGTGTCACCCTTAAGGAGAC<br>CAATAGAAACTAGGGCTCTGCGTTTGCTCCAGGTAGTCCGCTGCTCCCTTGGGCCTGGGCCCACTGACAGCCCTGGTG<br>CCTCTGGCCGGCTGCACACCTCCTGGCGGGCAGCTGTGAAAGTGCCGCCAGGTTTTGAGTGTTCTGGCAATACCTGCT<br>CAAAACCTGGCGGCACTTTCACGGAGGCCTGCCCTGACTGCCCACGGTGCCGTGGCCAAAGAGGATCTAAGGGCACC<br>GCTGAGGGCCTACCTAACCATCGTGGGGAATAAGGACAGTGTCACCCGGGCTTGTCGAGACAGAGAAGAGGGCTCTG<br>CGTTTGCTCCAGGTAGTCCGCTGCTCCCTTGGGCCTGGGCCCACTGACAGCCCTGGTGCCTCTGGCCGGCTGCACACCT<br>CCTGGCGGGCAGCTGTGAAAGTGCCGCCAGGTTTTGAGTGTTCTGGCAATACCTGCTCAAAACGAGCCGGCACTTTCA<br>CGGAGGCCTGCCCTGACTGCCCACGGTGCCGTGGCCAAAGAGGATCTAAGGGCACCGCTGAGGGCCTACCTAACCAT<br>CGTGGGGAATAAGGACAGTGTCACCCACTCTTGCGTTTCTGATAGGCACCTATTGGTCTTACTGACATCCACTTTGCCT<br>TTCTCTCCACAG | | | BrainVTA |

| | | | |
|---|---|---|---|
| **Subcloning of SpCas9-gRNAs to Noodles (Figures 2L and 3C)** | | | |
| $gRNA_{290}$-noodles | CACCGATCTTGCGGTACTCAAACTA | AAACTAGTTTGAGTACCGCAAGATC | Genewiz |
| $gRNA_{295}$-noodles | CACCGATCTTGGTGAGACTCAAATG | AAACCATTTGAGTCTCACCAAGATC | Genewiz |
| $gRNA_{292a\text{-}5p}$-noodles | CACCGCCAGCCTGTGATACTCAAAC | AAACGTTTGAGTATCACAGGCTGGC | Genewiz |
| $gRNA_{292a\text{-}3p}$-noodles | CACCGATCGGAAGAAAAGTGCCGCC | AAACGGCGGCACTTTTCTTCCGATC | Genewiz |
| **Subcloning of SpCas9-gRNAs response sequence to Noodles (Figures 2L and 3C)** | | | |
| $gRNA_{290}$-response | CGCGATCTTGCGGTACTCAAACTATGGTGCA | CCATAGTTTGAGTACCGCAAGAT | Genewiz |
| $gRNA_{295}$-response | CGCGATCTTGGTGAGACTCAAATGTGGTGCA | CCACATTTGAGTCTCACCAAGAT | Genewiz |
| $gRNA_{292a\text{-}5p}$-response | CGCGCCAGCCTGTGATACTCAAACTGGTGCA | CCAGTTTGAGTATCACAGGCTGG | Genewiz |
| $gRNA_{292a\text{-}3p}$-response | CGCGATCGGAAGAAAAGTGCCGCCAGGTGCA | CCTGGCGGCACTTTTCTTCCGAT | Genewiz |
| **Primers for amplification off-target binding sites** | | | |
| Chr2:63,286,171 | GGGTTTCAAAATCCCATGC | GGGAGAGATGTGAGCAAAGC | Genewiz |
| Chr3:20,596,888 | AATCCATGAGCATGGGAGAT | CTTCGGTGAAGAAGCTGGAT | Genewiz |
| Chr4:34,908,522 | AGCGCTTCAGAGTTCTGTCC | CCTGGCTGGTTATGGCTTT | Genewiz |
| Chr8:24,622,701 | AATGTTGATCCTGCCAATCC | GTCAATGGCCTTTCTCTACACA | Genewiz |
| chrX:49,430,534 | GGGAAACAACACCCTTCTCA | AATTTTGATGGGGATTGCAT | Genewiz |
| chrX:145,752,145 | GGGGCAACAACTGGAATGTA | AAACAAATGAGGGTGGGATG | Genewiz |
| Chr10:38,493,861 | TATGCTGATTTGGGGAAAAA | AAGCAGGCAAGTTCAGCATA | Genewiz |
| Chr12:110,194,354 | TCCCTTCCCCGCTATTTATC | ACTGCCAGCCAAGACTCATT | Genewiz |
| Chr15:5,483,510 | GCCAGGTATACGCTGTTTGA | TTCCTGACAATGGACCAAGTT | Genewiz |
| Chr3:66,757,564 | CTCGTCCCACCTCATTCATC | TGCTCAGGCCCATTAGATTT | Genewiz |
| Chr4:67,270,303 | TCATGAAGAAAGAGGTCTGCAC | CACGCACTCTTGAAATTCCTT | Genewiz |
| Chr6:120,484,589 | GCCTGCTGTCAGACAAACAC | TTTACCTGTGGAGCCACCTT | Genewiz |
| Chr8:62,551,743 | AGGCATTGACATGGGAAGAC | GCCCAAACAGAGATAACAGGTC | Genewiz |
| Chr8:96,645,474 | AGCTGTCACCAACAGGAAGG | TGGAATTGATCCATATCTTGTTTT | Genewiz |
| Chr9:40,261,442 | GCAAACATTCAGGAACAAAGG | TTCCACTTCAGAGTGACTTACCAG | Genewiz |
| chrX:164,652,010 | CACACTAAGCCAGCACATGG | ATTGTGGCCTTGAGCAAACT | Genewiz |

| | | | |
|---|---|---|---|
| Chr11:80,543,207 | CCCCTATTGCCCTGAGTGTA | TGCTGGGAATTGAACTTGGG | Genewiz |
| Chr11:82,315,768 | AAAGTTCCAGTCCTCCCCTCT | AGGTGAGCTTGAGGTGTGGT | Genewiz |
| Chr11:100,285,646 | GGTCACTGTAGAATGCCCCT | TGTTGTGAGGACTGAAGGCT | Genewiz |
| Chr12:29,213,836 | GTGGGGAGACCCACTATGTC | AGCCCCACACTGTAGTGACC | Genewiz |
| Chr15:101,201,595 | ATAAAGACACCACGGCCTGA | TTGGAAAGCCTAGGAGGTAGG | Genewiz |
| Chr2:36,070,359 | GCCCAGAGAATGAGGTGCTA | TTTGTCCCCTGCTTGAATGC | Genewiz |
| Chr2:153,537,552 | CACTGGGCTATGGGAAGTCA | GGGGAAATGGTGTGAGTCCT | Genewiz |
| Chr4:139,531,877 | CATGTCACCCCAACAACCAG | AAGGGAAGGAATGGAGGTGG | Genewiz |
| Chr17:3,777,103 | GCAGTGTTTAACGACTCCCA | TGATGGAGGCAGATCAGTGA | Genewiz |
| **Subcloning of SpCas9-gRNAs to double-gRNA cassette equipped rAAV vector** | | | |
| gRNA$_{290}$-rAAV | ACCGATCTTGCGGTACTCAAACTA | AACTAGTTTGAGTACCGCAAGATC | Genewiz |
| gRNA$_{295}$-rAAV | GATCTTGGTGAGACTCAAATGGTTTT | CATTTGAGTCTCACCAAGATCGGGAA | Genewiz |
| gRNA$_{292a\text{-}5p}$-rAAV | GCCAGCCTGTGATACTCAAACGTTTT | GTTTGAGTATCACAGGCTGGCGGGAA | Genewiz |
| gRNA$_{292a\text{-}3p}$-rAAV | ACCGATCGGAAGAAAAGTGCCGCC- | AACGGCGGCACTTTTCTTCCGATC | Genewiz |
| **Subcloning of SpCas9-gRNAs to one-gRNA cassette and Cas9 equipped HP180 vector** | | | |
| gRNA$_{290}$-HP180 | CACCGATCTTGCGGTACTCAAACTA | AAACTAGTTTGAGTACCGCAAGATC | Genewiz |
| gRNA$_{295}$-HP180 | CACCGATCTTGGTGAGACTCAAATG | AAACCATTTGAGTCTCACCAAGATC | Genewiz |
| gRNA$_{292a\text{-}5p}$-HP180 | CACCGCCAGCCTGTGATACTCAAAC | AAACGTTTGAGTATCACAGGCTGGC | Genewiz |
| gRNA$_{292a\text{-}3p}$-HP180 | CACCGATCGGAAGAAAAGTGCCGCC | AAACGGCGGCACTTTTCTTCCGATC | Genewiz |
| **Subcloning of SpCas9-gRNAs to double-gRNA cassette and Cas9 equipped HP180.2 vector (Figure 3A)** | | | |
| gRNA$_{292a\text{-}5p}$-HP180.2 | CACCGCCAGCCTGTGATACTCAAAC | AAACGTTTGAGTATCACAGGCTGGC | Genewiz |
| gRNA$_{292a\text{-}3p}$-HP180.2 | ACCGATCGGAAGAAAAGTGCCGCC | AACGGCGGCACTTTTCTTCCGATC | Genewiz |
| **In-fusion subcloning of Pten 3′UTR into the dual-luciferase vector pmirGLO (Figures 5E-H)** | | | |
| Pten-luc | AAACGAGCTCGCTAGCAGGGTTTTGACACTTG TTGTCCA | CGACTCTAGACTCGACTGGAGATGGT GTATGGTCCAGAG | Genewiz |

[†]DNA, RNA and locked nucleic acid (LNA) nucleotides are indicated with capital small and italic letters, respectively.

**Table S3. Off-target analysis of gRNAs.**

| gRNA ID | gRNA sequence[†] | Targeted gene[*] | Off-target binding site location[*] | Off-target sequence[#] | Off-target effiency of gRNA (%) | Potentially affected locus | Genomic context |
|---|---|---|---|---|---|---|---|
| $gRNA_{290}$ | ATCTTGC GGTACTC AAACTA*T GG* | miR-290 (Chromosome 7: 3,218,626-3,218,708) | Chr2:63,286,171 | gTCTTcCtGTACTCAAACTACGG | 1.9349 | Kch7 | intergenic |
| | | | Chr3:20,596,888 | CCTTAGTTaGAGTcaCGCAAGAT | 99.9399 | Gm31466 (lincRNA) | intergenic |
| | | | Chr4:34,908,522 | CCTTAcTTTGAGgACCGtAAGAT | 0.1821 | Platr9 | intronic |
| | | | Chr8:24,622,701 | CCTTAGTTaGATACacCAAGAT | 99.9855 | Adam18 | intronic |
| | | | chrX:49,430,534 | ATCTTGCGtgACTCtAACTAAGG | 99.9662 | Arhgap36, Gm14696 (lincRNA) | intergenic |
| | | | chrX:145,752,145 | CCCTAGTaTtAGTACaGCAAGAT | 0.306 | Gm15058 (processed pseudogene) | intergenic |
| $gRNA_{295}$ | ATCTTGG TGAGACT CAAATG*T GG* | miR-295 (Chromosome 7: 3,220,773-3,220,841) | Chr10:38,493,861 | ATCTTGGTGAtACTaAAATtTGG | 10.9198 | Gm48198 (processed pseudogene) | intergenic |
| | | | Chr12:110,194,354 | cCaTGGTGAGtCTCAAATGTGG | 0.3039 | Gm34785 (lincRNA) | intergenic |
| | | | Chr15:5,483,510 | tTCTTGGTGAGAaaCAAATGAGG | 0.4069 | Gm46496 (processed pseudogene) | intergenic |
| | | | Chr3:66,757,564 | CCCagTaTGAGTCTCACCAAGAT | 0.2549 | Gm6555 (processed pseudogene), Gm43516 (processed pseudogene), Rsrc1 | intergenic |
| | | | Chr4:67,270,303 | CCTCcaTTGAGTCTCACCAAcAT | 0.2243 | Gm11403 (processed pseudogene) | intergenic |

| | | | | | | | |
|---|---|---|---|---|---|---|---|
| | | | Chr6:120,484,589 | CCACATTTGAGTCTCAaCAgGAg | 0.2316 | Il17ra | exonic |
| | | | Chr8:62,551,743 | ATgTTtGTGAcACTCAAATGTGG | 0.3564 | Gm7561 (processed pseudogene) | intergenic |
| | | | Chr8:96,645,474 | CCGCATTTaAGTCTCACCtgGAT | 0.3147 | Gm24132 (miRNA) | intergenic |
| | | | Chr9:40,261,442 | AgCTTtcTGAGACTCAAATGAGG | 0.1692 | Scn3b | intergenic |
| | | | chrX:164,652,010 | ATCTTGaTGAatCTCAAATGAGG | 0.5859 | Gm23404 (snoRNA) | intergenic |
| gRNA$_{292a-5p}$ | CCAGCCTGTGATACTCAAAC*TGG* | miR-292a (Chromosome 7: 3,219,189-3,219,270) | Chr11:80,543,207 | CCAGTTgGAGcATCACAGcCTGG | 0.3779 | Myo1d | intronic |
| | | | Chr11:82,315,768 | aCAGgCTGTGATACaCAAACAGG | 0.421 | Ccl1, Tmem132e | intergenic |
| | | | Chr11:100,285,646 | CCAGTTTGgGcATCtCAGGCTGG | 1.2754 | Krt42, Eif1 | intergenic |
| | | | Chr12:29,213,836 | CCAGTgTGAGTtTCACAGGCTtG | 0.4764 | Gm31333, Myt1l | intergenic |
| | | | Chr15:101,201,595 | aCAGCCTGTGATgCTgAAACAGG | 0.3968 | Acvr1b | intronic |
| | | | Chr2:36,070,359 | CCTGTTTGtGTgTCAgAGGCTGG | 0.5677 | Morn5 | intronic |
| | | | Chr2:153,537,552 | CCAGCCTGgGAgACTCAAAgGGG | 3.1338 | Nol4l, Commd7 | intergenic |
| | | | Chr4:139,531,877 | CCAGCCTGccATAtTCAAACGGG | 0.4905 | Iffo2 | UTR5 |
| gRNA$_{292a-3p}$ | ATCGGAAGAAAAGTGCCGCC*AGG* | miR-292a (Chromosome 7: 3,219,189-3,219,270) | Chr17:3,777,103 | ATCGGgAGAAAtGTGCaGCCAGG | 1.1463 | Nox3, 4930470H14Rik | intergenic |

[†] PAM nucleotides (outlined by italic) are adjacent to the 3′end, but not included to the sequence.

[*] GRCm38.p6 mouse reference genome was used for analysis. Thresholds for mismatches of SpCas9 and SaCas9 gRNAs were set to 3 and 2, respectively.

[#] Mismatches are indicated by lowercase.

**Video S1.**
Free running activity and tail suspension assay in a 35-wk-old female miR-290-295$^{-/-}$ mouse.

**Video S2.**
Hindlimb clasping assay in 2-year-old females SN$^{DA}$:miR-290-295CKO and Control mice 55 wk after transduction.

**Video S3.**
Hindlimb clasping assay in 10-month-old males SN$^{DA}$:miR-290-295CKO$_{SaCas9}$, SN$^{DA}$:miR-290-295CKO$_{SaCas9}$-miR-292aOE and Control mice 20 wk after transduction.

**Video S4.**
Rotations in 10-month-old males SN$^{DA}$:miR-290-295CKO$_{SaCas9}$, SN$^{DA}$:miR-290-295CKO$_{SaCas9}$-miR-292aOE and Control mice 20 wk after transduction.

**Video S5.**
Rotations in 6-month-old females 6-OHDA, SN$^{DA}$:miR-292aOE-6-OHDA and Control mice 17 wk after transduction.
